# Microbiota – gut – brain axis modulation drives glioblastoma progression and therapy resistance

**DOI:** 10.64898/2026.05.14.724808

**Authors:** Sarah Lavielle, Gauthier Delrot, Maria M Haykal, Doriane Bomont, Marie-Alix Derieppe, Julie Martineau, Sébastian Lillo, Préscillia Alves Gomes, Erwan Guichoux, Corinne Buré, Benoît Pinson, Nathalie Dugot-Senant, Barbara Klink, Nathalie Nicot, Pol Hoffmann, Teresa Frisan, Ioannis S Pateras, Macha Nikolski, Thomas Daubon, Océane CB Martin

## Abstract

Glioblastoma is a highly aggressive brain tumour with poor prognosis, whose aetiology, progression, and therapeutic resistance remain incompletely understood. While the microbiota–gut–brain axis has emerged as a key regulator of neurological disorders, its role in glioblastoma biology and treatment response is still largely unexplored.

Using a clinically relevant immunocompetent murine model combining glioblastoma stem cell implantation, dextran sodium sulfate-induced gut inflammation, and a full Stupp-like therapeutic protocol, we investigated bidirectional gut–brain communication in glioblastoma. Tumour growth and recurrence were monitored by bioluminescence imaging, tumour transcriptomic profiles were analysed by RNA sequencing, and brain and colon tissues were subjected to histological and molecular analyses. Gut microbiota composition was assessed by 16S rRNA sequencing, while systemic metabolites and cytokines were quantified in plasma. Cross-compartment association bioinformatic analyses were performed to correlate multi-organ readouts.

Gut inflammation enhanced glioblastoma growth and promoted tumour recurrence following therapy. Tumour progression was associated with increased infiltration of immunosuppressive macrophages, whereas recurrence correlated with elevated oxidative DNA damage. Remarkably, glioblastoma exerted systemic immunomodulatory effects, attenuating intestinal and systemic inflammatory responses, and induced profound remodelling of gut microbiota composition and predicted metabolic function, including enrichment of *Akkermansia* and depletion of *Lactobacillus*. Systemic metabolic profiling was investigated as a route of communication within the gut-brain axis and revealed adaptations in DSS-treated mice associated with tumour burden and therapeutic response. Multi-compartment correlation and multivariable association analyses identified specific bacterial genera and circulating metabolites associated with tumour volume, intestinal inflammation, and genomic instability.

These findings uncover a dynamic, bidirectional microbiota–gut–brain axis in glioblastoma and identify intestinal inflammation as a critical determinant of tumour progression and therapeutic outcome. Targeting gut disturbances and microbiota-associated metabolic pathways may represent novel strategies to modulate glioblastoma aggressiveness and treatment response.

## Introduction

Glioblastoma is the most common and aggressive primary brain tumour in adults, characterized by a poor prognosis and a median survival of approximately 14 to 15 months.^1^ Despite the current clinical approach, known as the Stupp protocol, which includes surgical resection followed by radiotherapy and chemotherapy with temozolomide^2^, glioblastoma remains associated with high rates of recurrence and therapeutic failure. Therefore, improving patient survival and quality of life requires a deeper understanding of the factors involved in tumour initiation, progression, and treatment resistance.

Recently, the gut-brain axis has become a promising field of research to better understand brain function and neurological disorders. The bidirectional communication between the gut and the brain occurs via systemic circulation through metabolite exchange, neural pathways, and immune system modulation.^3^ The gut microbiota, a diverse community of microorganisms colonising the gastrointestinal tract^4^, is a key mediator of this communication. It plays a critical role in regulating brain functions^5^, including stress response^6^, neurogenesis^7^, myelination^8^, microglial maturation^9^, and blood-brain barrier integrity.^10^ Gut dysbiosis has been implicated in several neuropathologies, including Parkinson’s and Alzheimer’s diseases.^11^

Given the significant role of the gut in brain integrity, alterations in intestinal homeostasis are likely to influence the central nervous system. Notably, gut inflammation has been associated with increased anxiety and depression-like behaviours in mice.^12^ Inflammatory bowel disease (IBD) plays a significant role in neurological disorders and pathologies. For instance, patients suffering from IBD showed a higher risk of developing Parkinson’s disease and Multiple sclerosis.^13,14^ Additionally, chronic intestinal inflammation has been associated with cognitive dysfunction and behavioural changes in patients.^15^ Acute and chronic inflammation induced by dextran sulfate sodium (DSS) treatment in mice has been shown to increase neuroinflammation^16^ and affect neurogenesis.^17^ To date, few studies have investigated the role of intestinal inflammation in brain tumour development. In the context of medulloblastoma, DSS-induced gut inflammation in mice did not affect brain tumourigenesis.^18^ However, in patients, a Mendelian randomization study provided evidence supporting a potential causal link between IBD and brain cancer.^19^ Nonetheless, despite growing evidence linking gut inflammation to brain disorders, its role in glioblastoma progression and treatment resistance remains unexplored.

Here, we investigated the implication of intestinal inflammation on glioblastoma progression and recurrence using a DSS-induced colitis model.^20^ We show that intestinal inflammation enhances glioblastoma growth and promotes tumour recurrence following therapy. Conversely, the presence of glioblastoma and its treatment significantly alter intestinal physiology, leading to decreased intestinal inflammation and changes in microbiota composition. Moreover, DSS treatment induces systemic metabolic and inflammatory alterations associated with tumoral volume. Together, these findings demonstrate a bidirectional gut–brain communication axis in glioblastoma and identify intestinal inflammation as a previously underappreciated modulator of tumour aggressiveness and treatment outcome.

## Materials and Methods

### Animal experiments and samples

All mouse experiments and interventions were conducted in accordance with institutional guidelines and approved by the local and national ethics committee (agreement number: C33-522-22).

8-week-old C57BL/6N male mice were purchased from Charles River Laboratories. They were housed in pathogen-free conditions at the animal facility of Talence (Animalerie Mutualisée de Talence, University of Bordeaux, France) in rooms with controlled temperature, humidity, and light (22.0 ± 2.0 °C and 55.0 ± 10.0%, respectively; 12/12-hour light/dark cycle) and with *ad libitum* access to food and water.

A total of 60 mice were divided into six groups (n=10 per group): control mice (Ctl), DSS-treated control mice (DSS), mice implanted with murine glioblastoma stem cells (Impl), DSS-treated implanted mice (DSS Impl), mice undergoing the Stupp protocol (Stupp), and DSS-treated mice subjected to the Stupp protocol (DSS Stupp). The health status of mice was monitored throughout the experiment. Body weight was measured three times a week, and animals were euthanized if weight loss exceeded 20% of their maximal weight. In addition, the Disease Activity Index (DAI) was calculated, based on body weight loss, stool consistency, and the presence of fecal blood, as previously described.^21^

Each week, mouse faeces were harvested and stored at -20°C. Blood was collected by retro-orbital bleeding, prior to implantation and surgical resection, and after the complete Stupp protocol, and stored at -80°C. At the end of the experiment, mice were euthanized, and organs such as the brain, colon, and spleen were collected. Brain and colon were fixed in formalin and embedded in paraffin (FFPE) for further analysis.

### Cells

The mGB2 murine glioblastoma stem cell line was provided by Dr. Peter Angel^22^ from the German Cancer Research Center (DKFZ). Before implantation, they were cultured as spheres in Neurobasal medium (NBM) (Fisher Scientific, #11570556) supplemented with B27 (Fisher Scientific, #15360284), heparin (10 U/ml) (Sigma Aldrich, #H3149), 20 ng/mL bFGF (ThermoFisher Scientific, #100-18B) and Penicillin-Streptomycin (1,000 U/ml) (Dutscher, #P06-07100) and kept in a humidified 5% CO_2_ incubator at 37 °C. Cells were tested negative for mycoplasma by PCR with specific primers.

The cells were transduced with lentiviral particles (produced by the Vect’UB platform at University of Bordeaux) containing the MND-Luc-IRES2-ZSGreen plasmid, enabling luciferase expression.

### Tumour implantation and Stupp protocol treatment

Tumour implantation and Stupp protocol treatment followed a well-established procedure described by Pineau *et al*.^23^ Briefly, prior to the day of implantation, mGB2 cells were grown as spheroids.^24^ 10,000 mGB2 dissociated cells were seeded in NBM, 0.4% methylcellulose in a 96-well round-bottom plate. After three days of formation, five spheroids of mGB2 were intracranially implanted 2.2mm to the left of the Bregma and 3mm deep. The procedure took place under anaesthesia by isoflurane inhalation, preceded by 0.1mg/kg buprenorphine (i.p. injection) and local anaesthetic (Anesderm 5%). Mice received the Stupp protocol 28 days following implantation. Resection was performed under the same anaesthetic and analgesic procedure. Fluorescein was injected retro-orbitally to better visualize the tumour. A pump was used to remove the superior part of the tumour. After surgery, mice underwent three sessions of 2Gy radiotherapy, administered concomitantly with chemotherapy, and subsequently chemotherapy alone. The chemotherapeutic agent, temozolomide (Euromedex, #A-T1178) dissolved in 0.9% NaCl, was administered orally at a dose of 50mg/kg twice daily for 2 days, then once daily for 6 additional days.

Tumour development was monitored each week by bioluminescence imaging (Biospace).

### Fecal 16S DNA sequencing

Total DNA was extracted from faeces with QIAamp PowerFecal Pro DNA Kit (Qiagen, #51804) and sent to the PGTB platform of Bordeaux. Amplicon libraries were prepared according to the Illumina metagenomic sequencing library construction workflow, size-selected between 350 and 700 bp, and sequenced on an Illumina NextSeq 2000 (301|10|10|301 cycles configuration).

### Bulk tumour RNA sequencing

Tumours were dissociated using the Brain Tumor Dissociation Kit (Miltenyi, #130-095-942), adding a step of Debris Removal (Miltenyi, #130-109-398) and the gentleMACS™ Dissociator (Miltenyi). ZsGreen-positive cells were sorted by flow cytometry (BD FACsMelody^TM^ Cell Sorter) to ensure purity. RNA was isolated using the Direct Zol RNA Miniprep Kit (Zymo Research). DNase I treatment was performed on column during RNA purification. RNA quantity was assessed using an Implen Nanophotometer® N50 to measure A260/A230 and A260/A280 ratios, and a Qubit fluorometer (Thermo Fisher Scientific). RNA integrity was evaluated using a Fragment Analyzer System (Agilent) with the High Sensitivity RNA Kit, and both the Relative Quality Number (RQN) and DV200 values were determined. RQN values showed substantial variability between samples. Sequencing libraries were prepared using the SMART Seq® Stranded Kit (Takara), with a starting input of 1 ng of RNA. Libraries were quantified using the Qubit dsDNA HS Kit and fragment size distribution was assessed on the Fragment Analyzer System using the NGS Fragment Kit. All libraries were normalized prior to pooling and were sequenced on an Illumina NovaSeq 6000 platform to a target depth of 20 million paired-end reads per sample (2 × 75 bp).

### Gut histological scoring

Haematoxylin and eosin (H&E) staining of brain and colon samples was done by the histological platform of Bordeaux. 3-µm sections were obtained from paraffin-embedded tissue blocks. Sections were deparaffinized, rehydrated, stained with haematoxylin for 5 min, rinsed, and then counterstained with eosin for 3 min. Finally, the sections were dehydrated, cleared, mounted and imaged using a slide scanner (Nanozoomer). Histological severity of gut inflammation was assessed employing an additive score based on the separate evaluation of inflammatory cell infiltrate and intestinal architecture with a total score ranging from zero to six.^25^

### RNAscope

*In situ* mRNA labelling in the brain and colon was performed using RNAscope (RNAscope Multiplex Fluorescent Reagent Kit v2, ACD BioTechne, #323100) following manufacturer’s instructions. FFPE tissue slides were deparaffinized by immersion in toluene and absolute ethanol baths.

Target mRNAs *Il10* (# 317261), *Foxp3* (#432611), *Ifn-γ* (#311391), *Nos2* (#319131), *Il6* (#315891), as well as positive (#321811) and negative (#321831) control probes, were hybridized for two hours at 40°C using paired double-Z oligonucleotide probes supplied by BioTechne.

The fluorophore (TSA Vivid 520, 570 or 650) was coupled to the probe for 30 minutes at 40 °C. All fluorophores were diluted 1:1,500 except for TSA 650, diluted 1:3,000.

Nuclear counterstaining was performed with DAPI, and slides were mounted with ProLong Gold Antifade reagent (Thermo Fisher Scientific, #P36934). Slides were dried overnight and imaged on a Leica SP8 confocal microscope using a 63× objective. Stainings were analysed as previously described.^26^.

### Immunofluorescence

Immunostaining of 8-oxoguanine (8-oxoG) was performed on 6 µm FFPE sections of the brain and colon. Sections were deparaffinized in xylene and rehydrated through graded ethanol solutions. Antigen retrieval was carried out for 15 min at 95 °C in citrate buffer (10 mM sodium citrate, 0.05% Tween-20, pH 6). After washing in TBS/0.1% Tween, sections were permeabilized with 0.5% Triton X-100 and incubated with RNase A (100 µg/ml) for 1 h at 37 °C, followed by proteinase K for 10 min. DNA was denatured by incubation with 2 M HCl for 5 min, followed by neutralization with 1 M Tris-base for 7 min. After blocking with goat F(ab) anti-mouse IgG (1:100, 10% BSA), sections were incubated with anti-8-oxoG antibody (Abcam, #ab64548, 1:100). After washing in TBS/0.1% Tween, slides were incubated with the secondary antibody (AF647, 1:1,000, Invitrogen) for 1 h at room temperature and mounted with Vectashield mounting medium containing DAPI.

Analysis was performed using a custom ImageJ macro. The macro measures the mean fluorescence intensity per cell in both the nucleus and the cytoplasm, corresponding to the mitochondrial 8-oxoG signal, as previously described.^27^ For each sample slide (brain and colon), three to five 20X crops were analysed.

Immune cells staining was performed on 3-µm FFPE sections of brain and colon using a tyramide signal amplification (TSA) process (SAT704A001EA). Slides were deparaffinized and rehydrated, and antigen retrieval was performed in a 1 mM Tris–EDTA solution, pH 9. For brain sections, two primary antibodies were used: ARG1 (Cell Signaling, #93668S, 1:500) and IBA1 (Fisher Scientific, #13481357, 1:250). For colon sections, two primary antibodies were used: CD45 (Abcam, #ab10558, 1:500) and MPO (Fisher Scientific, #RB373A1, 1:200). Tissues were then incubated with the corresponding antibody for 45 min at room temperature. EnVision Flex/horseradish peroxidase (Dako-Agilent K800021-2) was used for signal amplification, revealed with the TSA kit coupled with FITC for IBA1 and MPO, and with the Cyanine 3 (Akoya Biosciences) kit for ARG1 and CD45. Nuclei were counterstained with Hoechst. Slides were imaged using a VS200 slide scanner (Olympus).

For all immunological markers, 3 to 5 10X crops were analysed using QuPath software, based on threshold-positive cell detection.

### Circulating inflammatory biomarkers

Systemic inflammation was assessed in plasma samples using ELISA kits according to the manufacturers’ instructions. IL-6 (Bio-Techne, #M6000B), lipocalin-2 (Bio-Techne, #DY1857) and IL-10 (Antibodies.com, #A323178) were measured using a CLARIOstar plate reader (BMG Labtech).

### Semi-targeted blood metabolomic analyses

Metabolites were extracted from blood (50µL) according to the protocol already described^28^ with the addition of heavy isotope internal standards used for data normalization: succinic acid-D_4_, L-aspartic acid-^13^C_4_,^15^N, L-tyrosine-^13^C_9_, and adenosine-D_14_ 5′-triphosphate. Samples were reconstituted in Milli-Q water (500 µL), insoluble material was removed by centrifugation (1 h, 4°C, 21,000 × g), and the supernatant was subjected to ultrafiltration using a Nanosep 10K Omega cartridge (Pall) for 15 min at 4°C and 14,000 × g.

Metabolites were separated by liquid chromatography using either high-performance ion chromatography (u-HPIC; ICS6000, Thermo Electron) or high-performance liquid chromatography (u-HPLC; Vanquish Flex, Thermo Electron), equipped with AS11-HC-4 µm (250 × 2 mm) and Acclaim RSLC PAII (2.2 µm, 120 Å, 2.1 × 150 mm) analytical columns, respectively. HPIC separations were performed at a flow rate of 0.38 mL/min with the discontinuous KOH gradient previously described.^29^ For HPLC, metabolites were resolved at 0.25 mL/min using mobile phases (A) 50 mM formic acid pH 3, and (B) methanol with a gradient from 0% to 90% (B) over 10 min followed by column re-equilibration.

For both chromatographic methods, metabolites were detected using a high-resolution Orbitrap mass spectrometer (Exploris 120, Thermo Fisher Scientific) equipped with an EASY-IC ion source operating in negative (HPIC and HPLC) and positive (HPLC) modes, with scan-to-scan lock-mass correction. Targeted metabolites were quantified using TraceFinder 5.2 SP3 (Thermo Fisher Scientific) from full-scan MS data (m/z 70–1,000) acquired at a resolution of 60,000, with data-dependent MS² scans acquired at 15,000 resolution with an HCD collision energy of 30%. Metabolite identification relied on retention time, accurate mass, natural isotopic distribution, and MS² fragmentation patterns.

### Bioinformatic analyses

#### Transcriptomics analysis

Raw FASTQ files were quality-checked and pre-processed with fastp^30^ (v1.0.1) to remove adapters and low-quality reads; QC reports were aggregated with MultiQC^31^ (v1.33.0). Reads were aligned to the mouse reference genome (GENCODE^32^ release M28; GRCm39) using STAR^33^ (v2.7.11b) with default parameters. Gene-level counts were obtained from the alignments (STAR option).

Differential expression analysis was performed in R using DESeq2^34^ (v1.42.0) on raw gene counts. Genes with an adjusted p-value (Benjamini–Hochberg correction) < 0.05 and a mean normalized count > 50 were considered significantly differentially expressed. Volcano plots were generated with EnhancedVolcano (https://github.com/kevinblighe/EnhancedVolcano) (v1.22.0).

Gene Ontology Biological Process^35^ enrichment analysis was performed with ClusterProfiler^36^ (v4.10.0) using org.Mm.eg.db (v3.18.0). Enrichment was tested against the set of all annotated genes as the background universe. Terms with adjusted p < 0.05 were considered significant. Gene annotations were retrieved with biomaRt^37^ (v2.58.0), and bubble plots were produced with ggplot2 (https://ggplot2.tidyverse.org) (v3.5.2).

#### Microbiome analysis

Paired-end sequences of 16S rRNA were obtained as FASTQ files. Quality control and adapter/low-quality trimming were performed using fastp (v1.0.1) and cutadapt^38^ (v4.2). Amplicon sequence variants (ASVs) were inferred with DADA2^39^ (v1.26.0), including error-model learning, dereplication, denoising, merging of paired reads, and chimera removal, yielding an ASV count table. Taxonomic assignment was performed against the SILVA^40^ database (release 138.1).

Downstream processing and visualization were carried out using phyloseq^41^ (v1.50.0) and microbiome (https://github.com/microbiome/microbiome) (v1.28.0). Differentially abundant taxa between DSS and control groups were identified using LEfSe^42^ (v1.1.2) with default parameters.

Then, to identify ASVs potentially associated with tumoral volume and gut inflammation, we conducted multivariable association testing using MaAsLin2^43^ (v1.16.0). MaAsLin2 was run using total sum scaling (TSS) normalization, log transformation, standardization and a prevalence filtering of 0.2; with tumoral volume and fecal lipocalin concentrations used as fixed effects. Associations were tested using a linear model and considered significant at q-value < 0.05 (Benjamini-Hochberg correction). Significant ASV-trait associations were visualized as a bipartite network, where nodes represent an ASV or a phenotypic trait, and edges represent significant associations. Edges were colored according to the effect size.

Functional profiling was inferred from 16S data using PICRUSt2^44^. Prior to inference, the ASV abundance matrix was TSS-normalized and filtered to retain ASVs present in at least 25 % of samples within each comparison. Inferred MetaCyc^45^ pathways were directly obtained from PICRUSt2 outputs., KEGG^46^ pathway abundances were computed from KO metagenomes using ggpicrust2 package^47^. Pathway abundance matrices (MetaCyc and KEGG) were log2 transformed and pairwise differential analysis was performed using non-parametrical Mann Whitney U test. Significantly different pathways were visualized as bar plots reporting the log2 ratio of group medians.

#### Semi-targeted blood metabolomic analyses

Metabolites with more than 25% missing values were excluded from the analysis. Remaining missing values were imputed using the metabolite-wise median across samples. Values were then Z-score normalised (mean-centred and scaled to unit variance). Volcano plots of differentially expressed metabolites were generated using MetaboAnalyst 6.0 (https://www.metaboanalyst.ca/).

For pathway-level analysis, KEGG metabolic pathways and their metabolites were downloaded using the KEGG API (release 117). Pathways represented by at least 3 metabolites in common with our dataset were retained. Pathway activity scores were computed using GSVA^48^ and the Gaussian kernel option for the empirical cumulative distribution function estimation. Differential pathway activity between condition pairs was assessed using the non-parametric Mann Whitney U test. Differentially active pathways were visualized using a clustered heatmap with Euclidean distance and average linkage. KEGG maps were generated using Pathview (v1.46.0)^49^ and mapped log2 fold changes using DIMet (v0.2.4)^50^.

#### Correlations with phenotypic traits

Pairwise associations between phenotypic traits were assessed in Python. Pearson correlation was used for pairs of continuous variables, and Spearman correlation was used when at least one of the variables was ordinal (DAI, Histological score). For continuous-continuous pairs, an ordinary least squares linear regression was additionally fitted to estimate the coefficient of determination (R²). P-values were computed using SciPy^51^, and associations were considered significant at *p* < 0.05.

### Statistical analysis

The data were graphed using GraphPad Prism 6. The experimental unit was defined as a single animal. First, normality was assessed, and statistical significance was determined using ANOVA (for three or more comparisons) or an unpaired Student’s t-test (for two-group comparisons). When needed, Welch’s correction or non-parametric equivalent was used. *p* < 0.05 was considered statistically significant.

## Results

### DSS treatment induced gut inflammation and changes in microbiota composition

To mimic the IBD condition characterized by chronic and cyclical intestinal inflammation, mice were given 1% dextran sulfate sodium (DSS) for six cycles of 5 days each (**Supplementary Fig. 1A**). Mice under DSS treatment exhibited transient weight loss after each cycle, followed by recovery during the subsequent periods (**Supplementary Fig. 1B).** The DSS treatment efficacy was validated by classical biological markers of colitis, such as a significant increase in fecal lipocalin levels, disease activity index (DAI), colon weight/length ratio, and spleen weight; and a significant decrease in the colon length in DSS-treated mice compared to control mice (**Supplementary Fig. 1C-G**). Histological evaluation verified the macroscopic findings, showing increased severity of inflammation in DSS-treated mice versus untreated ones (**Supplementary Fig. 1H and Fig. 3B-C**). DSS treatment also induced significant expected changes^52,53^ of the gut microbiota, such as an increase of *Escherichia-Shigella*, *Bacteroides*, *Romboustia*, *Turicibacter*, *Clostridium sensu stricto 1* and a decrease of *Alistipes*, *Actinobacteria*, *Eubacterium xulanophilum*, *Lactospiraceae_UGC_00* (**Supplementary Fig. 1I**).

### Gut inflammation enhances glioblastoma progression and recurrence

To investigate the impact of gut inflammation on glioblastoma progression and therapeutic resistance, mice, treated or not with DSS, received an intracranial injection of Luc^+^ mGB2 glioblastoma stem cells. Three weeks later, tumours in the Stupp and DSS-Stupp groups were surgically resected, followed by temozolomide administration and irradiation (**Fig. 1A**). Body weight was monitored throughout the procedure and did not differ significantly between groups (**Supplementary Fig. 2A**). Tumour burden was assessed by bioluminescence (**Fig. 1B**). Before the Stupp protocol, DSS-Impl mice harboured significantly larger tumours than non-DSS-implanted controls (**Fig. 1B bottom panel, 1C-D, Supplementary Fig. 2A**). Tumour regrowth after the Stupp protocol tended to be greater in DSS-treated mice, even though resection efficacy was similar between groups (**Fig. 1E**). Relapse occurred in five out of ten DSS-treated mice versus three out of ten non-DSS mice (**Fig. 1F**). Genomic instability induced by oxidative stress was assessed by 8-oxoguanine staining in tumours. The enhanced regrowth in DSS-Stupp mice was associated with increased mitochondrial and nuclear DNA damage within the tumour (**Fig. 1G-H**). In non-tumoral areas, 8-oxoG status remained unchanged in non-DSS-treated mice; however, DSS-treated mice exhibited elevated mitochondrial and nuclear 8-oxoG following tumour implantation, which was restored after the Stupp protocol (**Supplementary Fig. 2C**). These data indicate that gut inflammation is associated with enhanced glioblastoma growth and increased oxidative DNA damage following therapy.

**Figure 1:**
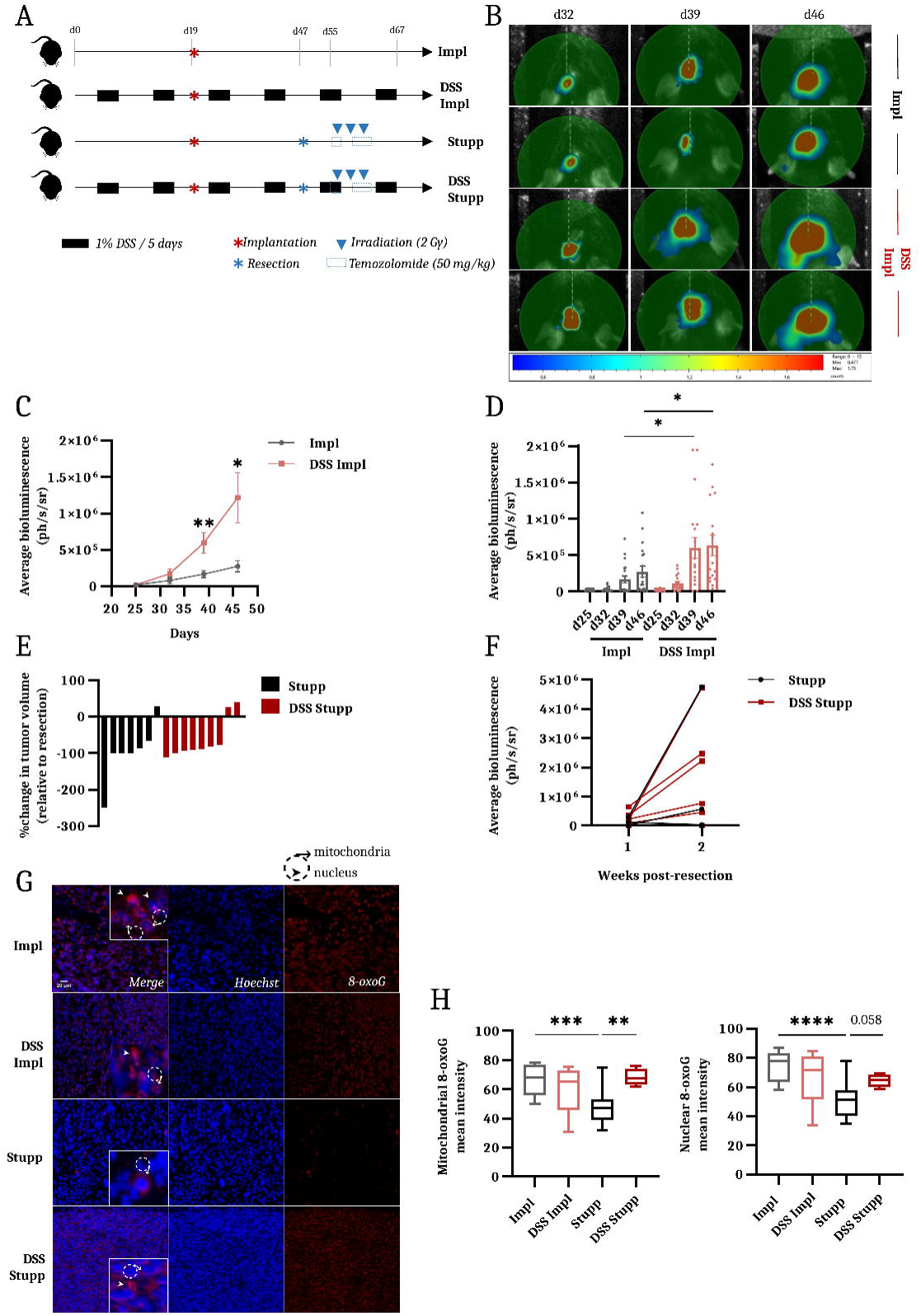
Gut inflammation accelerates glioblastoma progression and recurrence. **(A)** Experimental design of the study. Mice were pre-exposed or not to 1% DSS treatment cycles before intracranial implantation at day 19. Tumours were surgically resected at day 47, followed by radiotherapy (2 Gy) and temozolomide (50 mg/kg). Four experimental groups were analyzed: implanted control (Impl), DSS-treated implanted (DSS Impl), implanted Stupp-treated (Stupp), and DSS-treated Stupp-treated (DSS Stupp) mice. N = 10 mice per group. **(B)** Representative bioluminescence images of tumour-bearing mice from the Impl and DSS Impl groups at days 32, 39, and 46 post-implantation. Two mice per group are shown to illustrate tumour growth dynamics. **(C)** Tumour growth kinetics assessed as average bioluminescence in Impl and DSS Impl mice. Data are presented as mean ± SEM. **p < 0.01; *p < 0.05, two-way ANOVA. **(D)** Tumour bioluminescence values for all mice at each time point. Data are presented as individual values with mean ± SEM. *p < 0.05, unpaired t-test. **(E)** Waterfall plot showing the fold change in tumour volume a week after surgical resection relative to day 46 (the last pre-resection measurment). Each bar represents an individual mouse. **(F)** Tumour recurrence rates in Stupp and DSS Stupp groups following therapy. **(G)** Representative immunofluorescence images of brain sections stained for 8-oxoguanine (8-oxoG, red) to assess nuclear and mitochondrial oxidative DNA damage. Nuclei are counterstained with Hoechst (blue). Scale bar = 20 µm. **(H)** Quantification of mitochondrial (left panel) and nuclear (right panel) 8-oxoG signals in Impl, DSS Impl, Stupp, and DSS Stupp mice. Data are presented as median ± min/max. ****p < 0.0001; ***p < 0.001; **p < 0.01, unpaired t-test.

As macrophages represent the most abundant immune cells in glioblastoma^54^, we assessed the proportion of tumour-infiltrating macrophages, by *in situ* immunofluorescence. The overall proportion of IBA1^+^ macrophages was similar between the tumours of Impl and DSS-Impl mice; however, tumours of the DSS-Impl mice contained a significantly higher fraction of ARG1^+^ macrophages, indicating an immunosuppressive phenotype (**Fig. 2A-B, Supplementary Fig. 3A**). Consistently, RNAscope analysis revealed a significant reduction in *Nos2* expression in DSS-Impl tumours compared with Impl controls **(Fig. 2C-E, Supplementary Fig. 3B)**, supporting a shift toward macrophage-mediated immune suppression^55^. Following the Stupp protocol, both IBA1^+^ and ARG1^+^ macrophages increased significantly in tumours from non-DSS and DSS-treated mice (**Fig. 2A-B**). DSS-Stupp tumours displayed significantly reduced levels of *Il10* and *Ifn-γ* compared with Stupp controls, suggesting a global attenuation of immune signaling rather than a shift toward a defined inflammatory state (**Fig. 2C-E**). In contrast, *Nos2* expression was significantly increased in DSS-Stupp tumours compared to DSS-Impl tumours, supporting an aberrant inflammatory activation likely driven by therapy-induced stress (**Fig. 2C-E**). These findings support the emergence of a dysfunctional and immunologically silent tumour microenvironment in DSS-treated mice following therapy.

**Figure 2:**
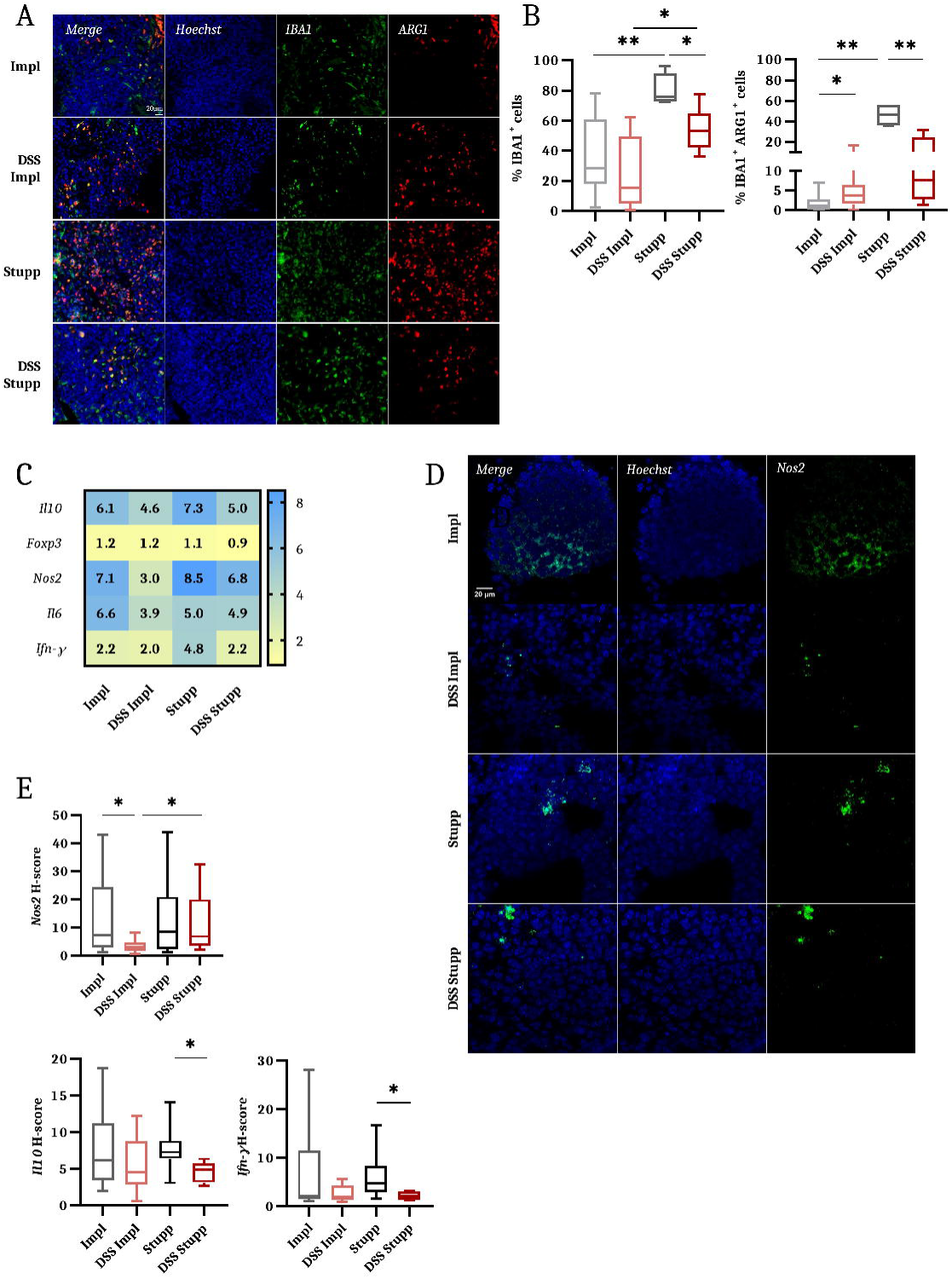
Gut inflammation reshapes the tumour immune microenvironment and promotes immunosuppressive macrophage polarization. **(A)** Representative immunofluorescence images of tumour sections from Impl, DSS Impl, Stupp, and DSS Stupp mice stained for IBA1 (green) to label macrophages and ARG1 (red) to identify immunosuppressive macrophages. Nuclei are counterstained with Hoechst (blue). Merged images are shown on the left and individual channels on the right. Scale bar = 20 μm. **(B)** Quantification of IBA1⁺ (left plot) and ARG1⁺ (right plot) cells expressed as a percentage of total cells in each experimental group. Data are presented as median ± min/max. **p < 0.01; *p < 0.05, unpaired t-test. **(C)** Heatmap summarizing RNAscope-based expression analysis of immune-related transcripts, including anti-inflammatory markers (*Il10*, *Foxp3*) and pro-inflammatory markers (*Nos2*, *Il6*, *Ifn-γ*), across Impl, DSS Impl, Stupp, and DSS Stupp tumours. Median H-score values are indicated. **(D)** Representative RNAscope images of tumour sections from Impl, DSS Impl, Stupp, and DSS Stupp mice using a *Nos2*-specific probe (green). Nuclei are counterstained with Hoechst (blue). Merged images are shown on the left and individual channels on the right. Scale bar = 20 μm. **(E)** Quantification of RNAscope signals expressed as H-scores for each transcript. Data are presented as median ± min/max. *p < 0.05, Mann-Whitney (*Nos2*, *Ifn-γ*), unpaired t-test (IL-10).

Beyond the tumour compartment, RNAscope analysis of non-tumoral brain regions revealed condition-specific immune alterations **(Supplementary Fig. 3C)**. DSS treatment alone altered brain immune signaling, with decreased *Il10* and increased *Nos2* expression, indicating that gut inflammation is sufficient to impact brain immune homeostasis. In non-DSS mice, glioblastoma implantation induced modest immunoregulatory changes in non-tumoral brain tissue, whereas DSS-treated mice exhibited a marked induction of *Il10* and *Foxp3*, consistent with immune preconditioning of the brain by gut inflammation **(Supplementary Fig. 3C**). Notably, the Stupp protocol disrupted this regulatory profile in DSS-treated mice, restoring *Nos2* expression and reducing *Il10* and *Foxp3* levels.

To further characterize the tumour-intrinsic response to therapy and determine whether it was influenced by intestinal inflammation, we performed transcriptomic profiling of Stupp-treated tumours. RNA sequencing revealed a marked upregulation of autophagy-related pathways in Stupp tumours (**Supplementary Fig. 3D-E**), reflecting an adaptive response to multimodal therapeutic stress. In contrast, DSS-Stupp tumours displayed reduced expression of gene sets associated with neuronal differentiation and synapse organization **(Supplementary Fig. 3F-G)**, indicating a loss of normal tissue identity and a microenvironment that may be increasingly permissive to tumour regrowth. This indicate that intestinal inflammation alters the molecular trajectory of tumours following therapy.

### Glioblastoma decreased gut and systemic inflammation

To investigate the bidirectional communication between the gut and the brain in the context of glioblastoma, we analysed the impact of tumour implantation and the Stupp protocol on intestinal physiology. In DSS-treated mice, the presence of glioblastoma was associated with reduced gut inflammation, as reflected by lower fecal lipocalin levels (**Fig. 3A**), a decreased colon weight/length ratio (**Supplementary Fig. 4A**), and a reduced severity of intestinal inflammation (**Fig. 3B-C**). This decrease in gut inflammation was reversed following the Stupp protocol, suggesting that the tumour itself transiently modulates colonic inflammatory tone.

**Figure 3:**
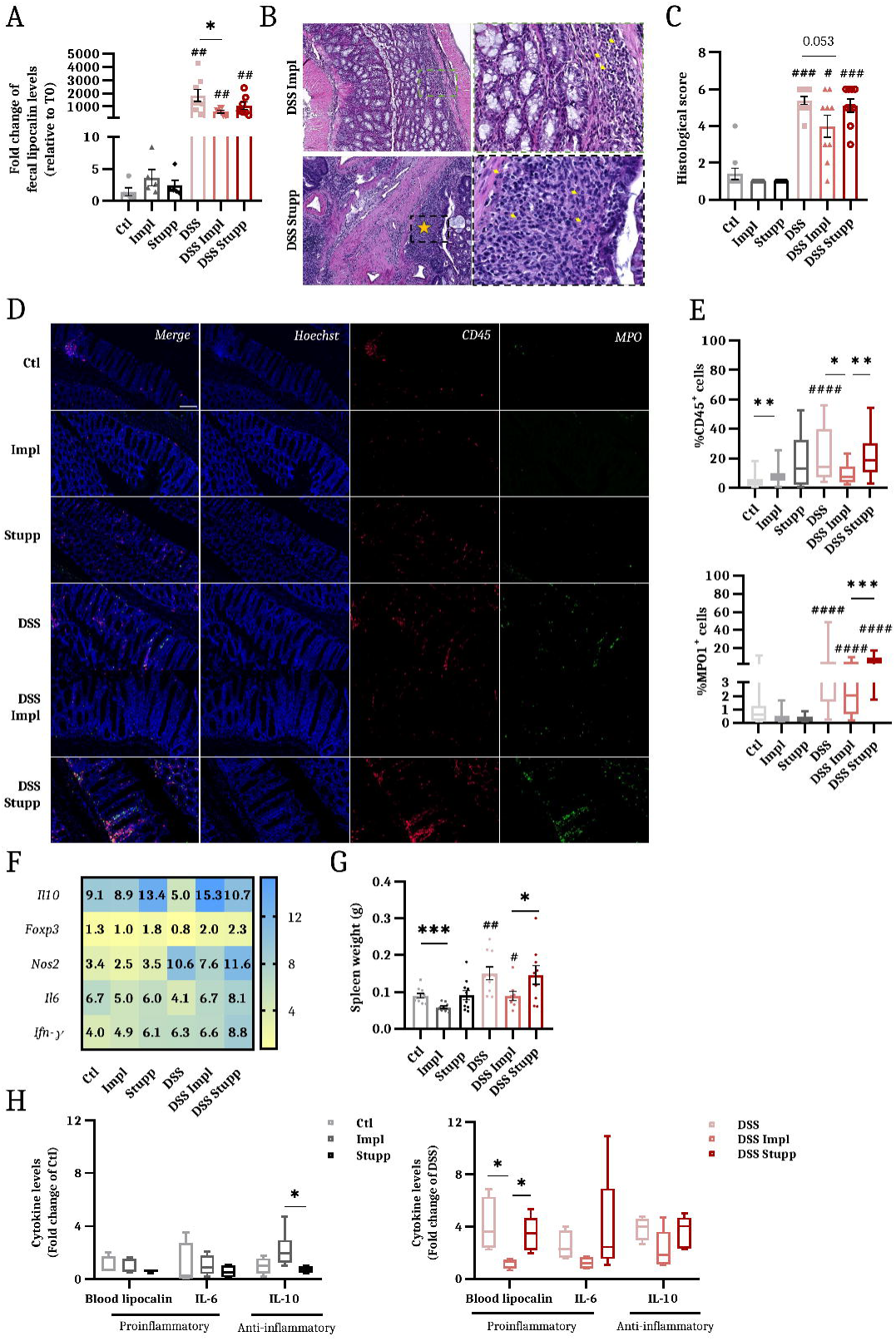
Glioblastoma attenuates intestinal and systemic inflammatory responses. **(A)** Fold change of fecal lipocalin levels measured at the end of the experiment relative to baseline. Data are presented as individual values with mean ± SEM. *p < 0.05; ##p < 0.01, Welch’s t-test. Hashtags indicate comparisons between Ctl *vs* DSS, Impl *vs* DSS Impl, and Stupp *vs* DSS Stupp groups. **(B)** Representative haematoxylin and eosin (H&E)-stained colon sections from DSS-treated Impl (top) and Stupp (bottom) mice. Low-magnification images (×20) are shown on the left, and high-magnification views (×400) on the right. Yellow arrows indicate neutrophil infiltration, and the asterisk marks areas of epithelial erosion. **(C)** Histological inflammation scores of colon sections across experimental groups. Data are presented as individual values with mean ± SEM. ###p < 0.001; #p < 0.05, unpaired t-test. Hashtags indicate comparisons between Ctl *vs* DSS, Impl *vs* DSS Impl, and Stupp *vs* DSS Stupp groups. **(D)** Representative immunofluorescence images of colon sections stained for CD45 (red) to label leukocytes and MPO (green) to identify neutrophils Nuclei are counterstained with Hoechst (blue). Merged images are shown on the left and individual channels on the right. Scale bar = 100 µm. **(E)** Quantification of CD45⁺ (left plot) and MPO⁺ (right plot) cells in the colon. Data are presented as median ± min/max. ***p < 0.001; **p < 0.01, *p < 0.05; ####p < 0.0001, Welch’s t-test. Hashtags indicate comparisons between Ctl *vs* DSS, Impl *vs* DSS Impl, and Stupp *vs* DSS Stupp groups. **(F)** Heatmap summarizing RNAscope-based expression analysis of immune-related transcripts, (*Il10*, *Foxp3*, *Nos2*, *Il6*, *Ifn-γ*) in colonic tissue across experimental groups. Median H-score values are indicated. **(G)** Spleen weight as a marker of systemic inflammation. Data are presented as individual values with mean ± SEM. ***p < 0.001; *p < 0.05, unpaired t-test; ##p < 0.01; #p < 0.05, Welch’s t-test. Hashtags indicate comparisons between Ctl *vs* DSS and Impl *vs* DSS Impl groups. **(H)** Plasma cytokine levels measured by ELISA in Ctl (left plot) and DSS-treated (right plot) groups. Data are presented as median ± min/max. *p < 0.05, unpaired t-test.

Tumour implantation exerted a marked distal effect on colonic homeostasis, affecting both oxidative stress and immune cell infiltration. In non-DSS mice, it increased mitochondrial and nuclear 8-oxoG in colonic cells, indicating elevated oxidative DNA damage that was partially restored following the Stupp protocol (**Supplementary Fig. 4B–C**). It was also associated with a 2-fold increase in CD45⁺ immune cell infiltration without significantly affecting MPO⁺ neutrophils (**Fig. 3D-E**), pointing to a reshaping of the colonic immune landscape rather than a neutrophil-driven inflammatory response. In contrast, in DSS-treated mice, where intestinal inflammation induced robust immune infiltration, tumour presence was associated with approximately a 2-fold reduction in CD45⁺ abundance and MPO⁺ cells, revealing a capacity of glioblastoma to dampen intestinal inflammation, an effect reversed by the Stupp protocol (**Fig. 3D–E**). RNAscope analysis of colonic immune signaling revealed condition-specific modulation of inflammatory markers (**Fig. 3F**). DSS treatment increased NOS2 and FOXP3 expression in the colon, in line with a concomitant activation of pro-inflammatory and compensatory regulatory responses in chronic intestinal inflammation. Glioblastoma implantation in DSS-treated mice was associated with a significant increase of IL10 and reduction of NOS2 expression, reflecting a shift toward a regulatory immune profile, which was partially disrupted following the Stupp protocol, with NOS2 significantly increasing in DSS-Stupp mice.

Glioblastoma also affected systemic inflammatory markers. Tumour-bearing mice exhibited reduced spleen weight, indicative of a lower systemic inflammatory state **(Fig. 3G)**. Circulating lipocalin was significantly decreased, and IL6 tended to be lower in DSS-implanted mice compared with DSS controls, reflecting a tumour-associated attenuation of systemic inflammation. Both parameters returned to control levels after the Stupp protocol, suggesting that therapy reverses these effects **(Fig. 3H**). Importantly, this effect appeared to be confined to DSS-treated mice with pronounced systemic inflammation, and may therefore not be observable under basal inflammatory conditions.

### Metabolomic profiling reveals systemic metabolic remodelling associated with gut inflammation and therapeutic response

To assess whether gut inflammation was associated with systemic metabolic alterations during glioblastoma progression and recurrence, we performed semi-targeted blood metabolomic profiling. Differential analysis revealed metabolic differences between experimental groups, as illustrated by volcano plot representations of DSS-regulated metabolites (**Fig. 4A, C**).

**Figure 4:**
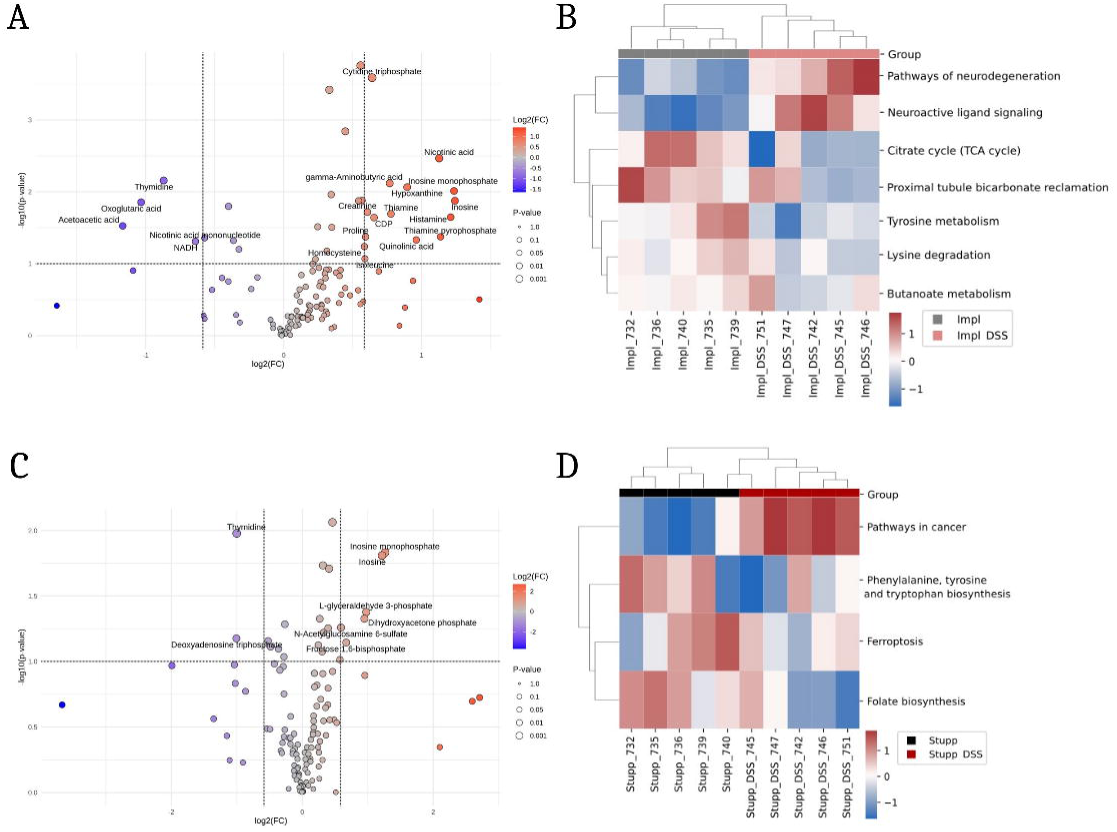
Gut inflammation drives systemic metabolic remodelling associated with tumour growth and therapeutic response. **(A)** Volcano plot depicting differentially abundant plasma metabolites between Impl and DSS Impl mice. Metabolites with a fold change ≥ 1.5 and *p* < 0.05 are highlighted as significantly upregulated (red) or downregulated (blue) in DSS Impl mice. **(B)** Heatmap of GSVA pathway scores derived from metabolomic profiling comparing Impl and DSS Impl mice, highlighting significantly altered metabolic pathways. **(C)** Volcano plot showing differentially abundant blood metabolites between Stupp and DSS Stupp mice. Metabolites with a fold change ≥ 1.5 and *p* < 0.05 are highlighted as significantly upregulated (red) or downregulated (blue) in DSS Stupp mice. **(D)** Heatmap of GSVA pathway scores derived from metabolomic profiling comparing Stupp and DSS Stupp mice, highlighting significantly altered metabolic pathways.

Comparison of DSS-Impl and Impl mice identified a distinct circulating metabolite profile associated with gut inflammation in tumour-bearing animals **(Fig. 4A)**. DSS-Impl mice exhibited increased levels of metabolites related to nucleotide metabolism and cellular stress, such as CTP, hypoxanthine, nicotinic acid, together with elevated neuroactive metabolites like gamma-aminobutyric acid (GABA) and histamine. In contrast, metabolites linked to energy metabolism and redox balance, were reduced, indicating a systemic metabolic shift associated with intestinal inflammation. Pathway-level analysis using z-score–based heatmaps revealed a coordinated downregulation of central carbon and energy metabolism pathways, including the tricarboxylic acid cycle, butanoate and tyrosine metabolism, alongside increased neuroactive ligand–receptor signaling **(Fig. 4B)**. To further illustrate the integration of differential metabolites within metabolic networks, key altered metabolites were mapped onto the KEGG purine metabolism pathway (**Supplementary Fig. 5**), highlighting a potential modulation of nucleotide turnover and salvage pathways in DSS-Impl mice.

Following therapy, DSS-Stupp mice displayed a different metabolic profile compared with Stupp controls, characterized by increased inosine and inosinic acid levels together with elevated glycolytic intermediates, while nucleotide-related metabolites such as deoxyadenosine triphosphate (dATP) and thymidine were reduced **(Fig. 4C)**. Pathway-level analysis revealed decreased activity of ferroptosis-related pathways and reduced folate and aromatic amino acid biosynthesis in DSS-Stupp mice **(Fig. 4D)**. Together, these data indicate that gut inflammation is associated with distinct, therapy-dependent systemic metabolic states accompanying enhanced tumour growth and recurrence, the mechanistic underpinnings of which warrant further investigation.

### Glioblastoma had an impact on gut microbiota composition

Given that circulating metabolites integrate host- and microbial-derived signals, we next investigated whether gut inflammation and tumour burden were associated with alterations in gut microbiota composition and with predicted metabolic functions.

Glioblastoma was associated with significant changes in gut microbiota of both untreated and DSS-treated mice (**Fig. 5A**). At the phylum level, GB-bearing mice showed a significant increase in *Verrucomicrobiota* and a decrease in *Firmicutes* (**Fig. 5B**). These changes were accompanied by a significant expansion of the *Akkermansiaceae* family (**Fig. 5C left panel)** and the *Akkermansia* genus **(Fig. 5D left panel),** irrespective of DSS treatment. Notably, both *Akkermansiaceae* and *Akkermansia* levels were restored following the Stupp protocol, indicating that therapeutic intervention partially normalized tumour-associated microbiota alterations. In parallel, glioblastoma implantation led to a significant reduction in *Lactobacillaceae* (**Fig. 5C right panel**) and *Lactobacillus* abundance **(Fig. 5D right panel)**, which remained only partially restored after therapy, particularly in non-DSS mice, suggesting a persistent alteration in specific bacterial taxa despite tumour resection and therapy.

**Figure 5:**
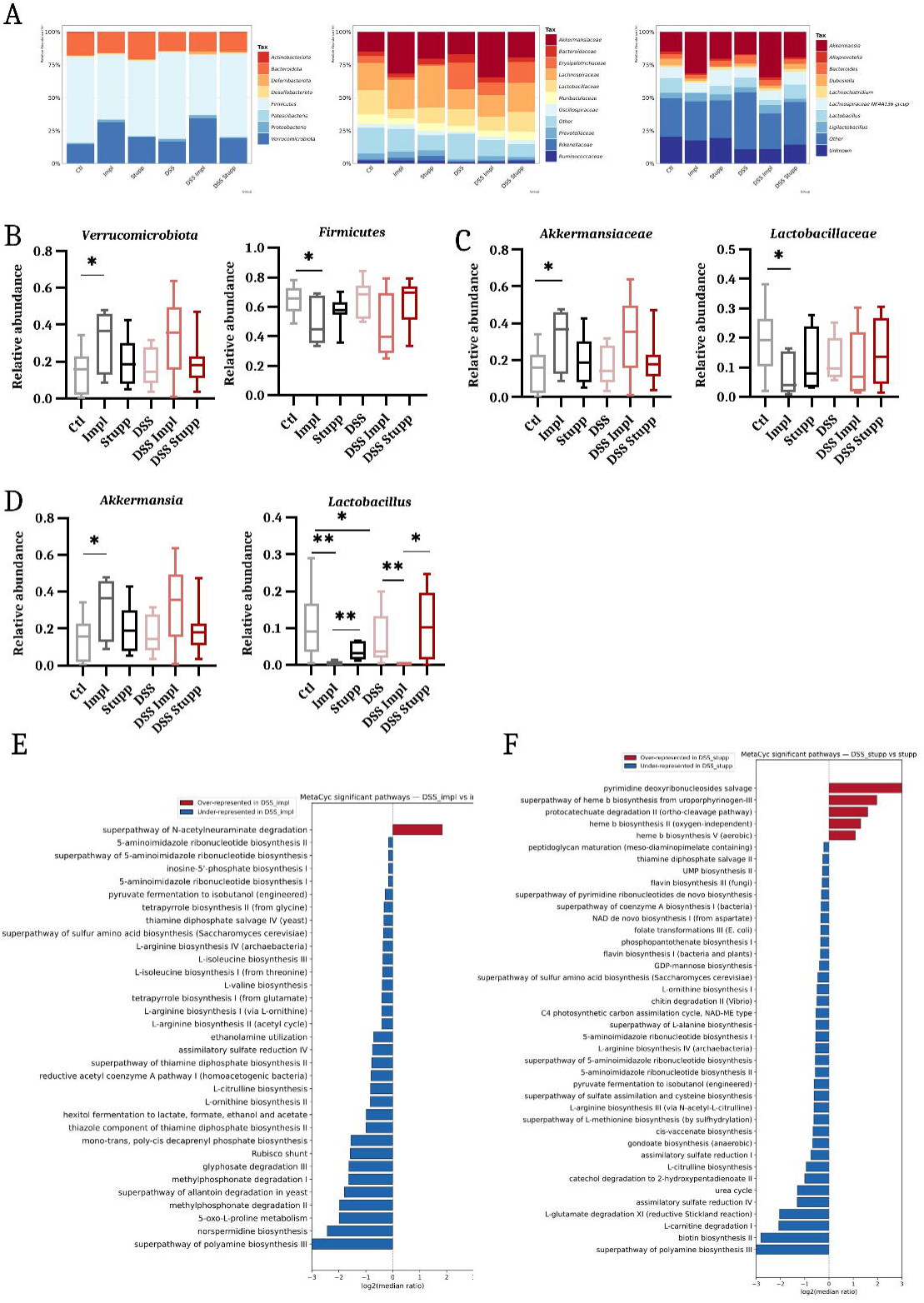
Glioblastoma reshapes gut microbiota composition. **(A)** Compositional bar plots of the gut microbiota across experimental groups at three taxonomic levels: phylum (left), family (middle), and genus (right). Relative abundances are displayed, and values represent group means. **(B)** Relative abundance of the phyla *Verrucomicrobiota* (left plot) and *Firmicutes* (right plot) across experimental groups. Data are presented as median ± min/max. *p < 0.05, unpaired t-test. **(C)** Relative abundance of the families *Akkermansiaceae* (left plot) and *Lactobacillaceae* (right plot) across experimental groups. Data are shown as median ± min/max. *p < 0.05, unpaired t-test. **(D)** Relative abundance of the genera *Akkermansia* (left plot) and *Lactobacillus* (right plot) across experimental groups. Data are presented as median ± min/max. **p <0.01; *p < 0.05, unpaired t-test. **(E-F)** Bar plot representation of significantly differentially abundant inferred functional pathways between Impl *vs* DSS Impl **(E)** and Stupp *vs* DSS Stupp **(F)**, annotated against the MetaCyc database. Pathways enriched in DSS Impl or DSS Stupp mice are shown in red, whereas pathways decreased relative to their respective controls are shown in blue. Functional predictions were derived from 16S rRNA gene sequencing data. Values correspond to the log₂ ratio of group medians.

To explore the functional consequences of glioblastoma implantation on gut microbial metabolism, we conducted metagenome functional prediction using PICRUSt with MetaCyc and KEGG. Tumour-bearing mice showed predicted enrichment in pathways related to secondary metabolite production, carotenoid biosynthesis, and anaerobic energy metabolism, alongside increased glycosaminoglycan degradation, while pathways involved in amino acid biosynthesis, peptidoglycan and heme production, bile acid metabolism, and carbohydrate utilization were predicted to decrease **(Supplementary Fig. 6A)**.

In DSS-treated, tumour-bearing mice, inflammation was associated with further predicted reductions in amino acid and polyamine biosynthesis, sulfate assimilation, and certain fermentative pathways, while sialic acid degradation was predicted to increase **(Fig. 5E, Supplementary Fig. 6B)**. Following the Stupp protocol, DSS-treated mice showed predicted persistent suppression of amino acid, polyamine, and energy metabolism pathways, together with increased nucleotide salvage, heme biosynthesis, and stress-response pathways **(Fig. 5F, Supplementary Fig. 6C)**. Notably, some of these predicted alterations parallel the systemic metabolic perturbations observed in blood metabolomics, including changes in nucleotide, energy, and amino acid metabolism; however, whether these reflect a functional microbiota-host metabolic axis or independent responses to the same biological stressors remains to be established.

### Gut bacterial traits were associated with tumoral volume

To integrate the multi-organ readouts collected throughout the study, we performed multivariable association analysis and statistical correlations, linking gut inflammation, microbiota composition, systemic metabolism, and tumour-associated parameters. Multivariable analysis revealed that tumour volume was significantly associated with several genera of the *Lachnospiraceae* family (FCS020 group 9, *Tyzzerella*, and *Eubacterium xylanophilum*), as well as *Ruminococcus*, *Anaerotruncus*, *Turicibacter*, and *Oscillospiraceae Intestinimonas* (**Fig. 6A**). Notably, *Tyzzerella* and *Turicibacter* were also positively associated with fecal lipocalin, identifying these taxa as potential microbial nodes linking tumour burden and intestinal inflammation (**Fig. 6A**).

**Figure 6:**
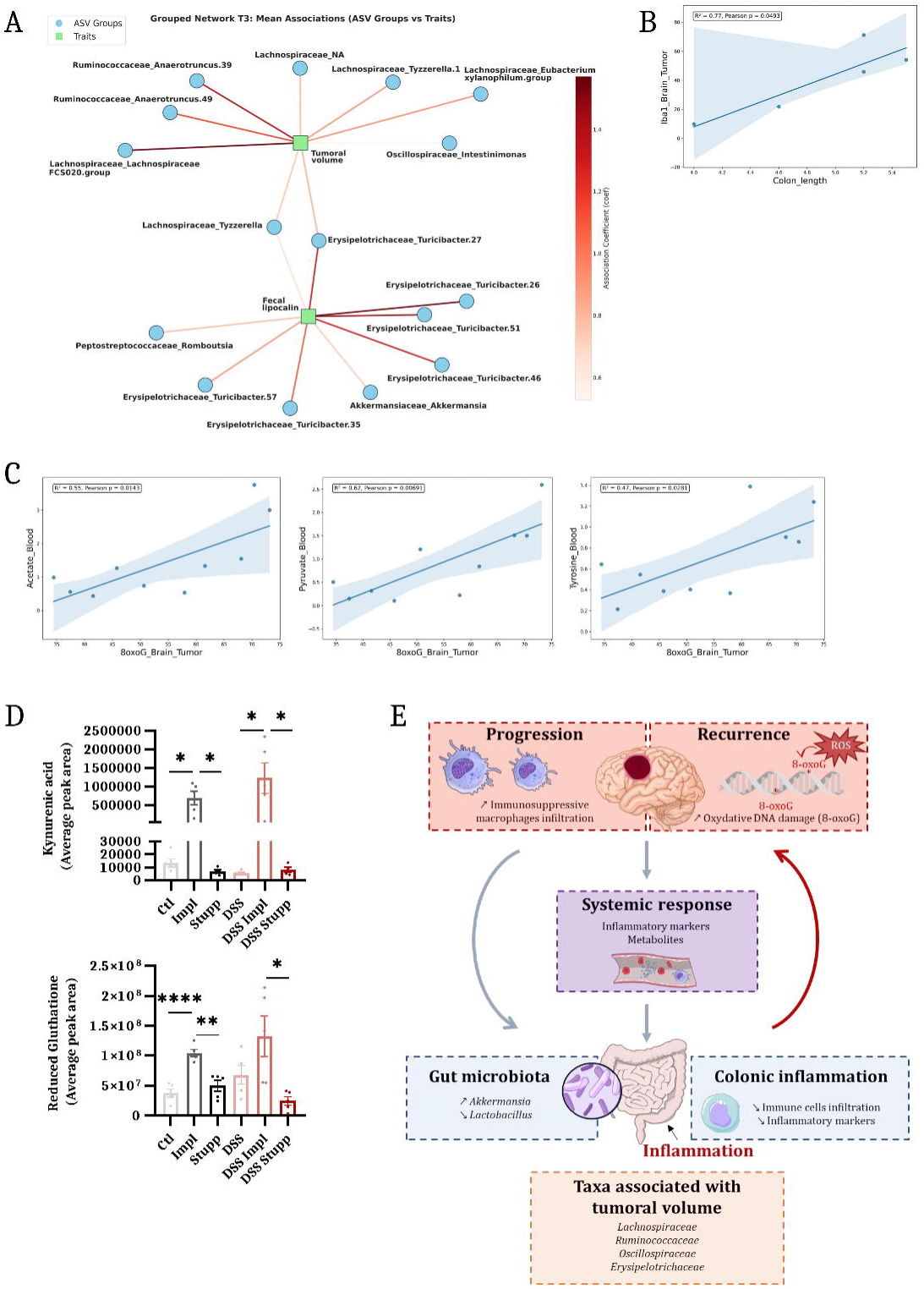
Multilevel associations linking gut inflammation, microbiota, systemic metabolism and glioblastoma progression. **(A)** Bipartite network representation of multivariable association analysis performed using MaAslin2. Nodes represent biological entities (phenotypic traits or ASVs), and edges indicate statistically significant associations. Edge colour intensity reflects the strength of the association. **(B)** Scatter plot illustrating the relationship between colon length and tumour-associated IBA1⁺ macrophage infiltration. The solid line represents the linear regression model. The corresponding *p*-value and R² are indicated. **(C)** Scatter plots showing correlations between tumour oxidative DNA damage (8-oxoG statuts) and circulating metabolites, including acetate (left plot), pyruvate (middle plot), and tyrosine (right plot). The solid lines indicate linear regression models, with corresponding p-values and R² displayed. **(D)** Average peak area of circulating kynurenic acid (upper plot) and reduced glutathione (bottom plot) across experimental groups. Data are presented as individual values with mean ± SEM. ****p < 0.0001; **p < 0.01; *p < 0.05, Welch’s t-test; Mann-Whitney. **(E) Intestinal inflammation promotes glioblastoma progression and recurrence within a bidirectional gut–systemic–brain axis.** DSS-induced intestinal inflammation enhances glioblastoma growth and is associated with an increased proportion of immunosuppressive ARG1⁺ macrophages in tumours. Following the Stupp protocol, intestinal inflammation is linked to increased tumour recurrence and elevated oxidative DNA damage (8-oxoG). Conversely, glioblastoma modulates peripheral physiology by reducing colonic immune cell infiltration and inflammatory marker expression and by reshaping systemic inflammatory and metabolic profiles. Tumour presence is also associated with gut microbiota remodelling, characterized by increased *Akkermansia* and decreased *Lactobacillus*. In addition, several bacterial taxa, including members of the *Lachnospiraceae*, *Ruminococcaceae*, *Oscillospiraceae* and *Erysipelotrichaceae* families, show positive associations with tumour volume.

Correlation analyses further uncovered significant relationships between gut and tumour immune parameters. In DSS-treated mice, tumour-associated macrophage abundance (IBA1⁺) positively correlated with colon length (**Fig. 6B**), indicating that intestinal inflammation was associated with reduced macrophage infiltration within the tumour microenvironment. Systemic metabolite profiling further revealed robust associations between tumour oxidative DNA damage and circulating metabolites. Tumour 8-oxoG status positively correlated with blood pyruvate, acetate, and tyrosine concentrations (**Fig. 6C**), linking systemic metabolic remodelling to tumour genomic instability. Finally, circulating kynurenic acid and reduced glutathione levels (**Fig. 6D**) were increased in tumour-bearing mice and further elevated in DSS-treated animals. These findings point to the activation of antioxidant defences and the tryptophan – kynurenine pathway, two central axes involved in immune regulation, oxidative stress adaptation, and tumour progression.

Together, these integrative analyses uncover coordinated microbiota – gut – systemic – tumour interaction networks and identify microbial and metabolic signatures associated with glioblastoma progression and intestinal inflammation.

## Discussion

The implication of the microbiota-gut-brain axis is well documented in neurological disorders^56^ and is emerging as a critical component of neuro-oncology^57,58^. Here, we provide the first comprehensive demonstration that intestinal inflammation modulates glioblastoma progression and therapeutic outcome. By integrating a clinically relevant glioblastoma stem cell model with a full Stupp-like treatment protocol, we show that gut inflammation enhances tumour growth, promotes recurrence after therapy, reshapes the tumour immune microenvironment, and reprograms systemic metabolism and gut microbial composition. Collectively, our findings support a model in which glioblastoma progression is embedded within a dynamic, bidirectional gut–microbiota–brain network (**Fig. 6E**).

### Gut inflammation promotes glioblastoma progression through immune and metabolic remodelling

A central finding of our study is that DSS-induced intestinal inflammation significantly enhances glioblastoma growth. in line with epidemiological evidence linking IBD to increased cancer incidence and mortality.^59^ Moreover, systemic inflammatory markers have recently emerged as predictors of poor outcome in glioma patients^60^, further supporting a functional link between inflammation and glioblastoma aggressiveness. In line with these observations, DSS-treated mice exhibited a robust systemic inflammatory response, characterized by a fourfold increase in circulating lipocalin and a 2.5-fold elevation in plasma IL6 levels, supporting the presence of a sustained systemic inflammatory state.

At the tumour level, DSS treatment profoundly reshaped the immune microenvironment. Glioblastoma is classically characterized by a highly immunosuppressive microenvironment, in which tumour-associated macrophages represent up to 50% of the cellular content^61^, consistent with the approximately half of tumour cells being IBA1⁺ observed in both groups. While total macrophage infiltration remained unchanged, DSS-treated tumours exhibited a significant enrichment in ARG1⁺ immunosuppressive macrophages accompanied by reduced *Nos2* and *Ifn-γ* expression, in line with tumour-promoting myeloid polarization known to facilitate immune evasion and invasion.^62,63^

In parallel, transcriptomic and metabolomic analyses revealed metabolic remodelling within tumours and systemically from DSS-treated mice. RNA sequencing uncovered a downregulation of glucose metabolism, redox homeostasis, and RNA processing pathways, while metabolomics revealed reduced metabolites linked to energy production and redox balance, including NADH, oxoglutarate, acetoacetate, and thymidine. Concomitantly, metabolites involved in nucleotide synthesis, cellular stress responses, and neuromodulatory signaling, such as inosine, hypoxanthine, nicotinic acid, and GABA, were increased. These changes suggest a systemic metabolic shift associated with intestinal inflammation, in line with emerging models of metabolic plasticity in glioblastoma, and warrant direct mechanistic interrogation in future studies.^64,65^

### Intestinal inflammation exacerbates therapeutic resistance and tumour recurrence

Importantly, gut inflammation also impacted therapeutic outcome. DSS-treated mice subjected to the Stupp protocol exhibited a trend toward increased tumour recurrence, accompanied by alterations in immune and oxidative stress parameters. As previously reported^66^, the Stupp protocol increased tumour-associated macrophage infiltration in non-DSS mice. However, in DSS-treated animals, this response was profoundly altered, as DSS-Stupp tumours displayed reduced IBA1⁺ and ARG1⁺ macrophage populations together with lower *Il10* and *Ifn-γ* expression. This immune profile indicates a global attenuation of immune activity rather than a shift toward a defined inflammatory state, supporting the emergence of a dysfunctional and immunologically silent tumour microenvironment.

Strikingly, this immune alteration was accompanied by increased oxidative DNA damage. Oxidative stress is a known driver of genomic instability, therapeutic resistance, and clonal evolution in glioblastoma.^67^ In this context, DSS-induced intestinal inflammation may exacerbate therapy-induced oxidative stress, fostering the emergence of aggressive tumour subclones and thereby promoting recurrence. Transcriptomic signatures of reduced neuronal differentiation and synaptic organization further support a shift toward immature, stem-like tumour states, which are closely associated with radioresistance and relapse.

### Glioblastoma exerts systemic and intestinal immunomodulatory effects

In line with the established capacity of solid tumours to reshape systemic immunity^68^, glioblastoma was associated with significant modulation of intestinal and systemic inflammatory states. In DSS-treated mice, tumour implantation attenuated colonic inflammation, as reflected by reduced fecal lipocalin, immune cell infiltration, and circulating inflammatory markers, suggesting that glioblastoma induces systemic immunoregulatory mechanisms capable of dampening peripheral inflammatory responses.

Glioblastoma is well known to promote systemic immunosuppression through circulating cytokines, myeloid-derived suppressor cells, and lymphocyte dysfunction.^69^ Our results extend this concept to the gut, indicating that intracranial tumours remotely modulate intestinal immune homeostasis. This systemic immunoregulatory effect may involve tumour-derived soluble mediators, metabolic reprogramming, or microbiota-driven pathways, and may represent an adaptive strategy to restrain excessive inflammation while preserving tumour-supportive immune niches.

### Glioblastoma-associated gut dysbiosis and functional remodelling

Along these systemic effects, glioblastoma reshaped gut microbiota composition and predicted functional capacity. Tumour-bearing mice exhibited enrichment of *Akkermansia* and depletion of *Lactobacillus*, together with a shift toward microbial pathways associated with anaerobic energy metabolism, stress adaptation, and reduced biosynthetic capacity. Similar alterations have been reported in glioblastoma patients, albeit with substantial inter-individual variability, supporting the translational relevance of our findings.^70–72^

Notably, several bacterial genera belonging to the *Lachnospiraceae* family, as well as *Turicibacter*, *Anaerotruncus*, and *Intestinimonas*, were positively associated with tumour volume. These taxa are known modulators of host immune tone and metabolic homeostasis, notably through the production of short-chain fatty acids and other bioactive metabolites, whose effects are highly context-dependent and may promote either immune tolerance or inflammation depending on the host and disease setting.^73–76^ Although causal relationships cannot be inferred from correlation analyses, these associations suggest that microbiota-mediated immune and metabolic signals may contribute to glioblastoma progression. The partial restoration of microbial composition following therapy further indicates that tumour burden is a major driver of dysbiosis, while treatment-induced stress imposes additional selective pressures on the microbial ecosystem.

### An integrated microbiota-gut-systemic-brain network in glioblastoma

Overall, our data uncover a coordinated bidirectional interplay between gut inflammation, microbiota remodelling, systemic immunometabolic reprogramming, and tumour adaptation that governs glioblastoma growth and recurrence (**Fig. 6E**). Intestinal inflammation reshapes tumour immunity and metabolism, while the tumour itself reprograms intestinal immune responses, microbial composition, and systemic metabolic homeostasis.

From a clinical perspective, our findings raise the possibility that intestinal inflammatory status and microbiota composition could represent modifiable determinants of glioblastoma outcome. Therapeutic strategies aimed at restoring gut immune and microbial homeostasis, such as dietary interventions or microbiota-targeted therapies, may hold promise as adjuvant treatments to improve therapeutic efficacy and limit recurrence.

### Limitations, conclusion, and further work

This study relies on a murine model of DSS-induced colitis, which does not fully recapitulate the complexity of human inflammatory bowel diseases. Moreover, while our integrative analyses reveal strong associations across biological compartments, causal mechanisms remain to be elucidated. Future studies with fully paired sampling across omics layers and increased cohort sizes will enable more mechanistic, predictive integration of microbiota–metabolite–host pathways. Finally, validation in patient samples will be essential to determine the translational relevance of these findings.

## Data availability

All datasets (16S rRNA gene sequencing, RNA-seq, and metabolomics datasets) generated in this study have been deposited in the European Nucleotide Archive (ENA) under accession number PRJEB111369.

## Acknowledgements

The author would like to thank Dr. Jean Descarpentrie and Manon Lemaitre for their technical help.

The authors thank the staff of the SAM Metabolic Analyses, Vect’UB, and FACSility platforms at TBMCore (University of Bordeaux, CNRS UAR 3427, INSERM US05) as well as the PGTB for their valuable assistance and expertise in the acquisition and analysis of data.

## Supplementary material

**Supplementary Figure 1: DSS treatment induces chronic intestinal inflammation and gut microbiota alterations**

**(A)** Schematic representation of the experimental timeline. Control (Ctl, top) and DSS-treated (bottom) mice received either regular water or 1% DSS for 5 days followed by 2 days off, over 6 cycles at days 0, 11, 25, 39, 53, and 64. n = 10 per group.

**(B)** Body weight monitoring of Ctl and DSS-treated mice over the experimental period. Body weight was measured twice weekly and is represented as mean ± SEM. DSS cycles are indicated as grey sections on the plot. ** p < 0.01; * p <0.05, unpaired t-test.

**(C)** Fold change of fecal lipocalin levels in Ctl and DSS-treated mice at the end of the experiment relative to the baseline. Data are presented as individual values with mean ± SEM. **** p < 0.0001, unpaired t-test.

**(D)** Disease Activity Index (DAI) in Ctl and DSS-treated mice at the end of the experiment relative to the baseline. Data are presented as individual values with mean ± SEM. **** p < 0.0001, unpaired t-test.

**(E)** Colon length in Ctl and DSS-treated mice. Data are presented as individual values with mean ± SEM. *** p < 0.001, unpaired t-test.

**(F)** Ratio of colon weight to colon length in Ctl and DSS-treated mice. Data are presented as individual values with mean ± SEM. **** p < 0.0001, unpaired t-test.

**(G)** Spleen weight in Ctl and DSS-treated mice. Data are presented as individual values with mean ± SEM ** p < 0.01, unpaired t-test.

**(H)** Representative haematoxylin and eosin (H&E)–stained sections of colons from Ctl (top) and DSS-treated (bottom) mice. Images on the left show low magnification (×20), and those on the right show high magnification (×400). Yellow arrows indicate infiltration of neutrophils; asterisk denotes an area with extended erosion.

**(I)** LDA score plot from LEfSe analysis identifying differentially abundant taxa between Ctl and DSS-treated mice at the end of the experiment.

**Supplementary Figure 2: Tumour growth dynamics and oxidative DNA damage in non-tumoral brain tissue**

**(A)** Body weight monitoring of of tumour-bearing mice throughout the experiment. Left panel: Impl and DSS Impl groups. Right panel: Stupp and DSS Stupp groups. Data are presented as mean ± SEM. Each DSS cycle is indicated as a grey section on the plot, implantation by a straight line, and the Stupp protocol by a non-continuous line.

**(B)** Waterfall plot showing the change in tumour volume in Impl and DSS Impl mice at day 46 relative to the first measurement after implantation. Each bar represents an individual mouse.

**(C)** Quantification of mitochondrial (left panel) and nuclear (right panel) 8-oxoG status in non-tumoral brain regions across all six experimental groups. Data are presented as median ± min/max. ** p < 0.01; * p < 0.05, unpaired t-test.

**Supplementary Figure 3: Tumour immune microenvironment characterization and transcriptomic profiling**

**(A)** Scatter plot illustrating the relationship between and tumour-associated IBA1⁺ and ARG1^+^ macrophage infiltration. The solid line represents the linear regression model. The corresponding *p*-value and R² are indicated.

**(B)** Representative RNAscope control images showing negative (top) and positive (bottom) controls. Sections were stained with the POLR2A probe (green). Nuclei are counterstained with Hoechst (blue). Merged images are shown on the left.

**(C)** Heatmap summarizing RNAscope-based expression analysis of immune-related transcripts, (*Il10*, *Foxp3*, *Nos2*, *Il6*, *Ifn-γ*), across experimental groups in non-tumoral brain tissue. Mean H-score values are indicated.

**(D)** Volcano plots showing differentially expressed genes in Stupp *vs* Impl tumours identified by RNA-seq analysis. Genes with a log₂ fold change > 1 and p < 0.05 are highlighted in red.

**(E)** Bubble plot showing functional enrichment analysis in Stupp *vs* Impl tumours. Bubble colour represents the adjusted p-value, bubble size indicates the number of genes (count), and the x-axis represents the gene ratio.

**(F)** Volcano plots showing differentially expressed genes in Stupp *vs* DSS Stupp tumours identified by RNA-seq analysis. Genes with a log₂ fold change > 1 and p < 0.05 are highlighted in red.

**(G)** Bubble plot showing functional enrichment analysis in Stupp *vs* DSS Stupp tumours. Bubble colour represents the adjusted p-value, bubble size indicates the number of genes (count), and the x-axis represents the gene ratio.

**Supplementary Figure 4: Glioblastoma-associated oxidative stress and immune remodelling in colonic tissue**

**(A)** Ratio of colon weight to colon length in DSS-treated, Impl, and Stupp mice. Data are presented as individual values with mean ± SEM. *p < 0.05, unpaired t-test.

**(B)** Representative immunofluorescence images of colon sections stained for 8-oxoG (red) assessing nuclear and mitochondrial oxidative DNA damage. Nuclei are counterstained with Hoechst (blue). Zoomed fields are shown in the right corner of the merged channels. Scale bar = 50 µm.

**(C)** Quantification of mitochondrial (top panel) and nuclear (bottom panel) 8-oxoG status in colonic regions across all six experimental groups. Data are presented as median ± min/max. * p < 0.05; ##p < 0.01; #p < 0.05, unpaired t-test. Hashtags indicate comparisons between Ctl *vs* DSS and Impl *vs* DSS Impl.

**Supplementary Figure 5: Mapping of differentially abundant circulating metabolites onto the KEGG purine metabolism pathway**. Data were integrated into the KEGG purine metabolism pathway to visualize the metabolic context of altered nucleotide-related metabolites in tumour-bearing mice exposed to gut inflammation. Metabolites detected in the dataset are highlighted in yellow, while differentially abundant metabolites between DSS-Impl and Impl mice are indicated in red (upregulated) or blue (downregulated).

**Supplementary Figure 6: Glioblastoma and gut inflammation alter predicted gut microbial functional capacity**

**(A)** Bar plot representation of significantly differentially abundant inferred functional pathways between Impl *vs* Ctl, annotated against the MetaCyc (left plot) and KEGG (right plot) databases. Functional predictions were derived from 16S rRNA gene sequencing data. Values correspond to the log₂ ratio of group medians.

**(B-C)** Bar plot representation of significantly differentially abundant inferred functional pathways between DSS Impl *vs* Impl **(B)** and DSS Stupp *vs* Stupp **(C)**, annotated against the KEGG database. Functional predictions were derived from 16S rRNA gene sequencing data. Values correspond to the log₂ ratio of group medians.

