## Supplementary Figures for "Microbiota – gut – brain axis modulation drives glioblastoma progression and therapy resistance"

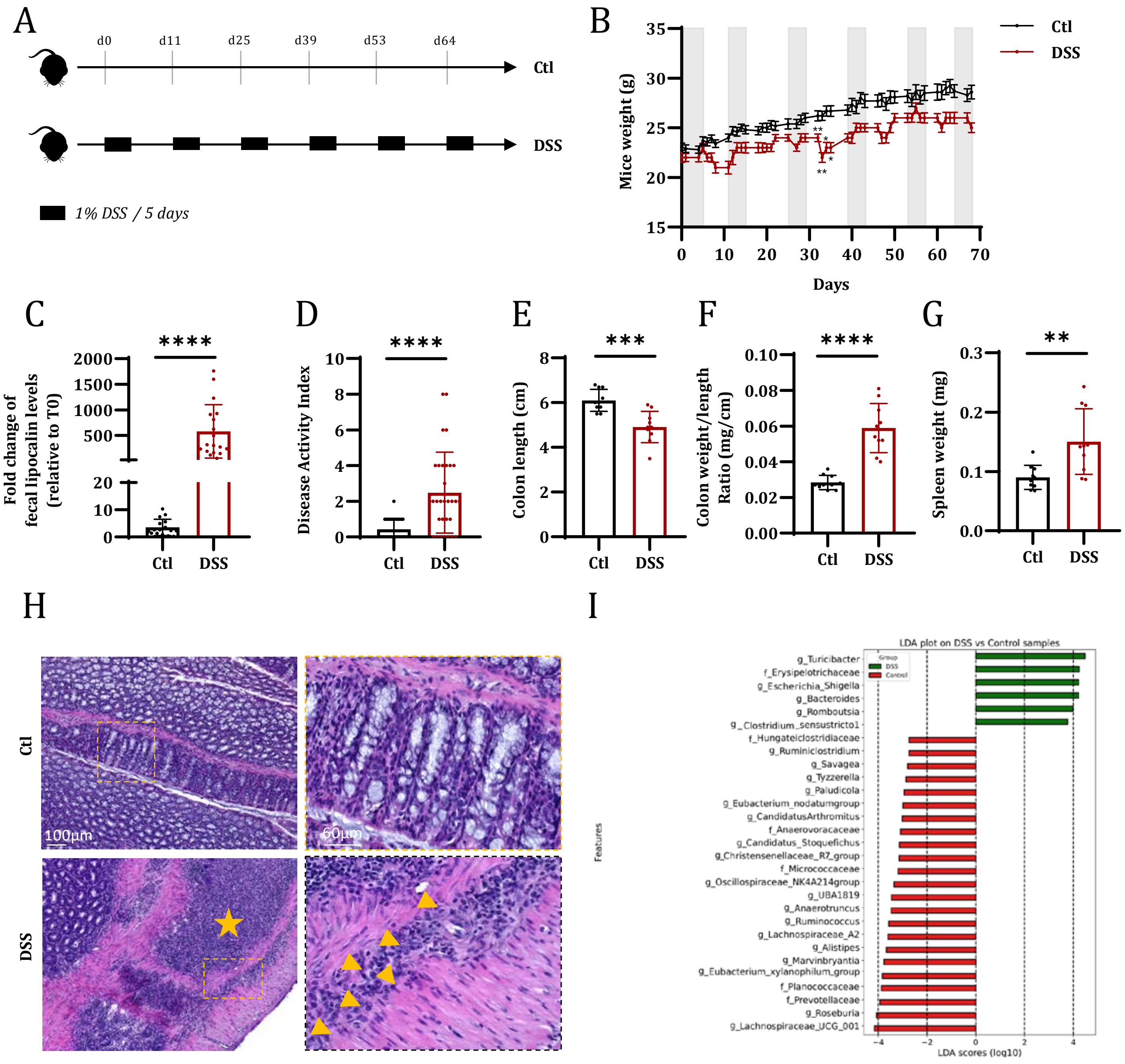


**Supplementary Figure 1**

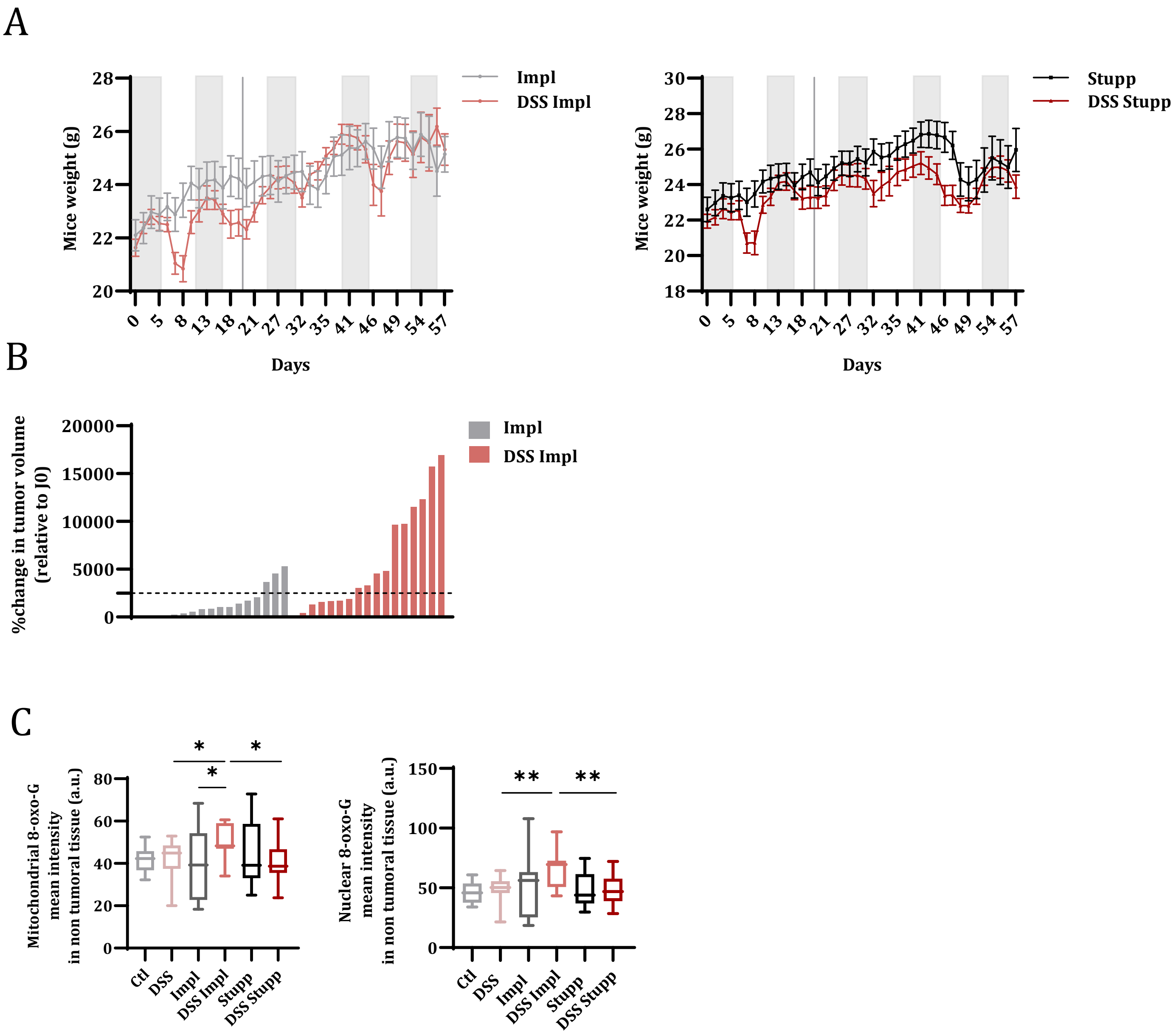


Supplementary Figure 2

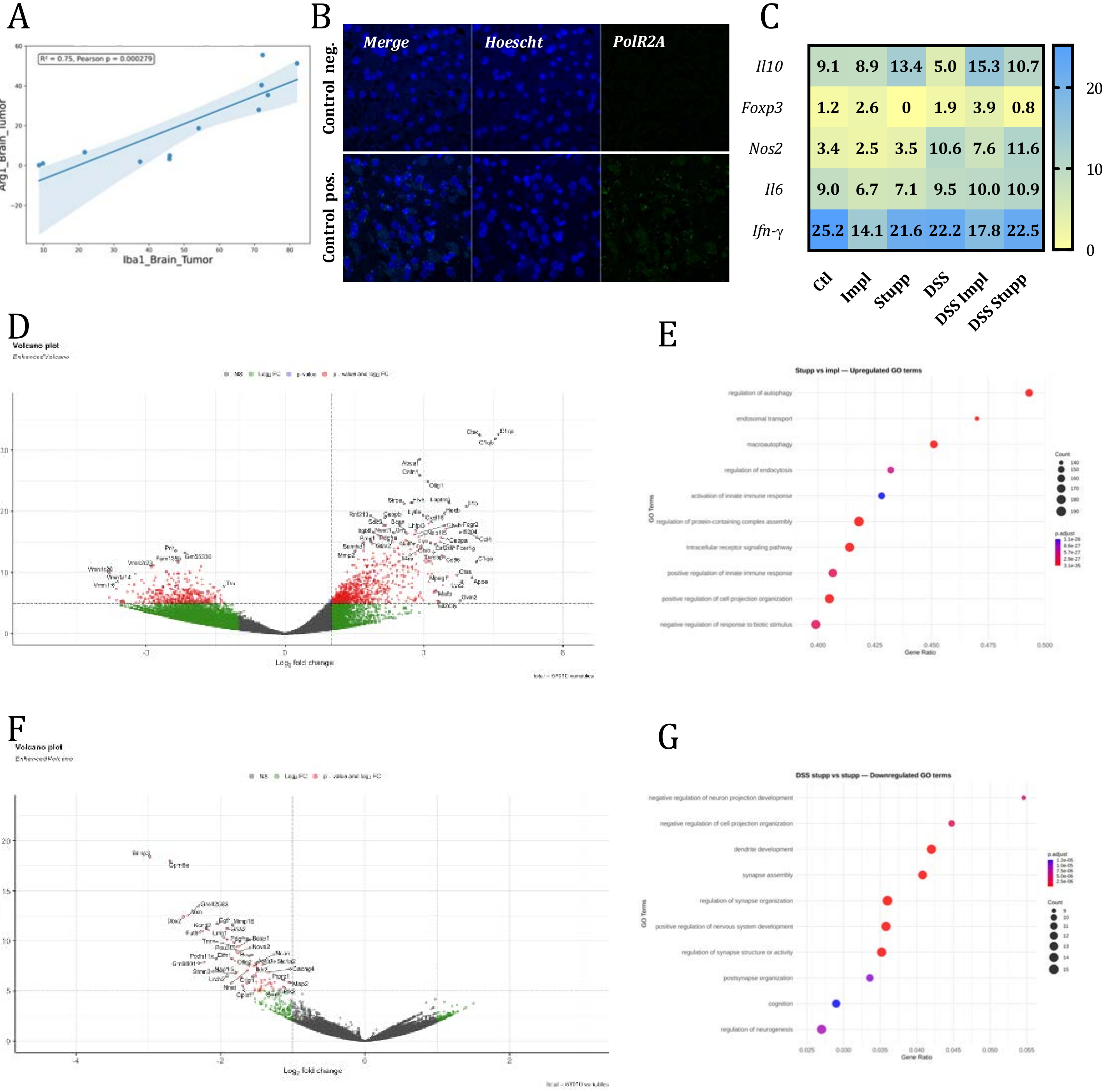


**Supplementary Figure 3**

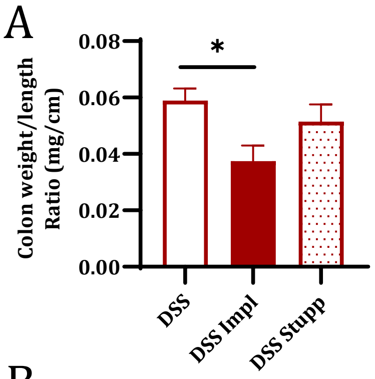


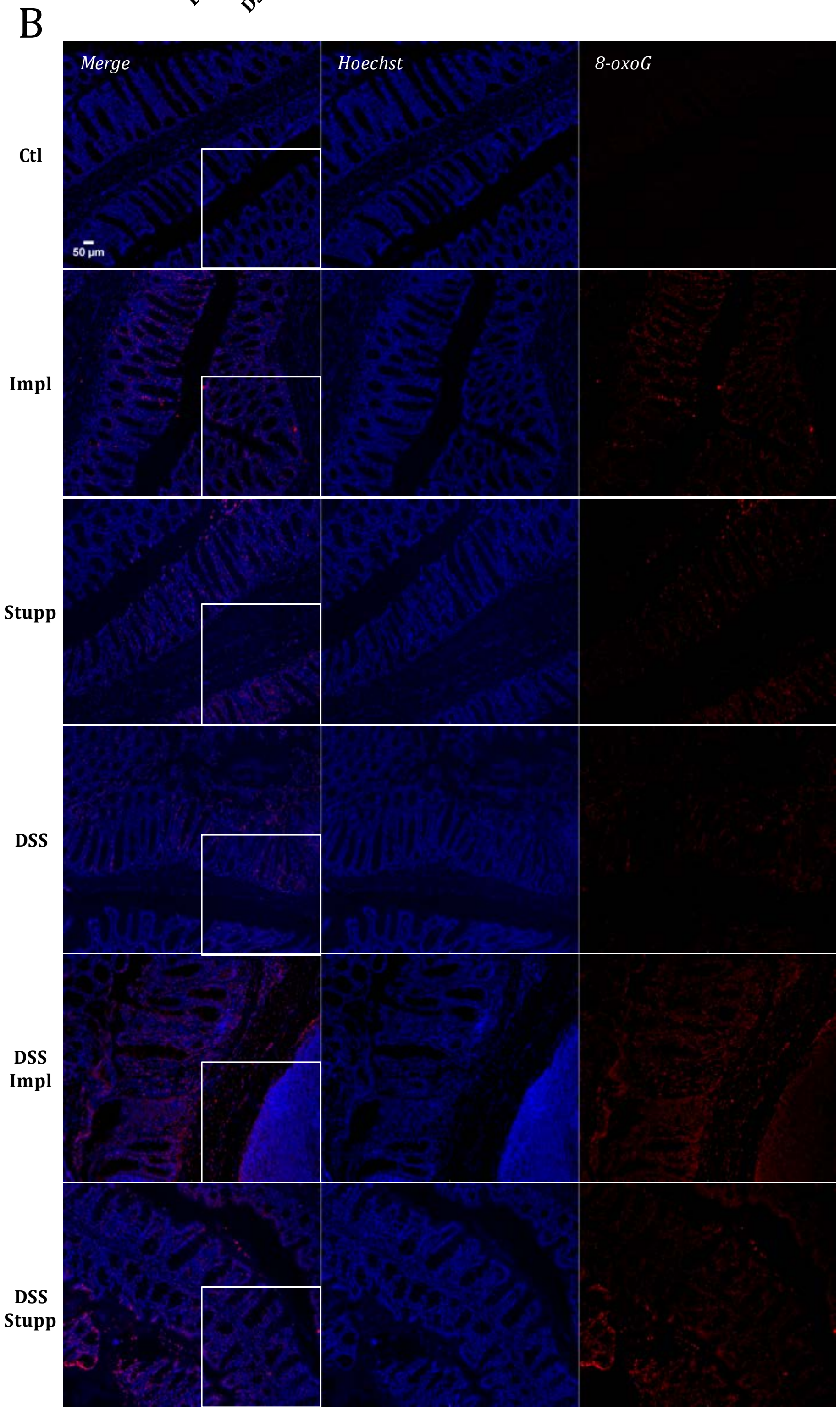


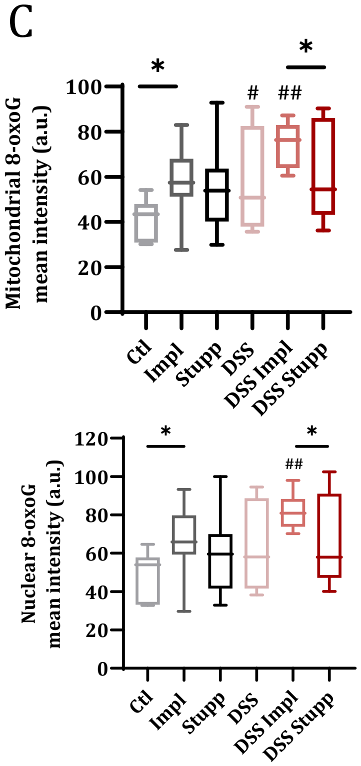


**Supplementary Figure 4**

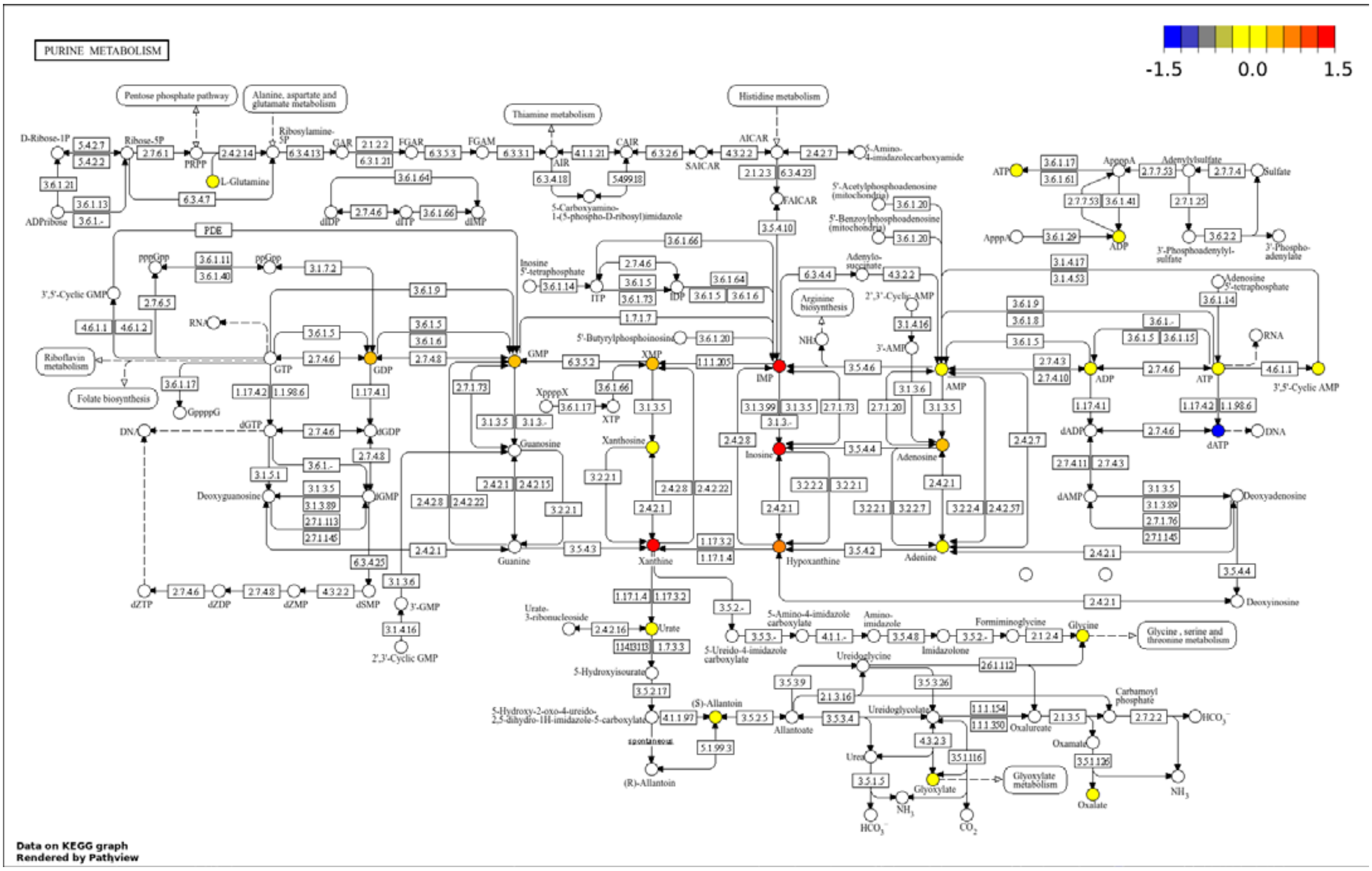


**Supplementary Figure 5**

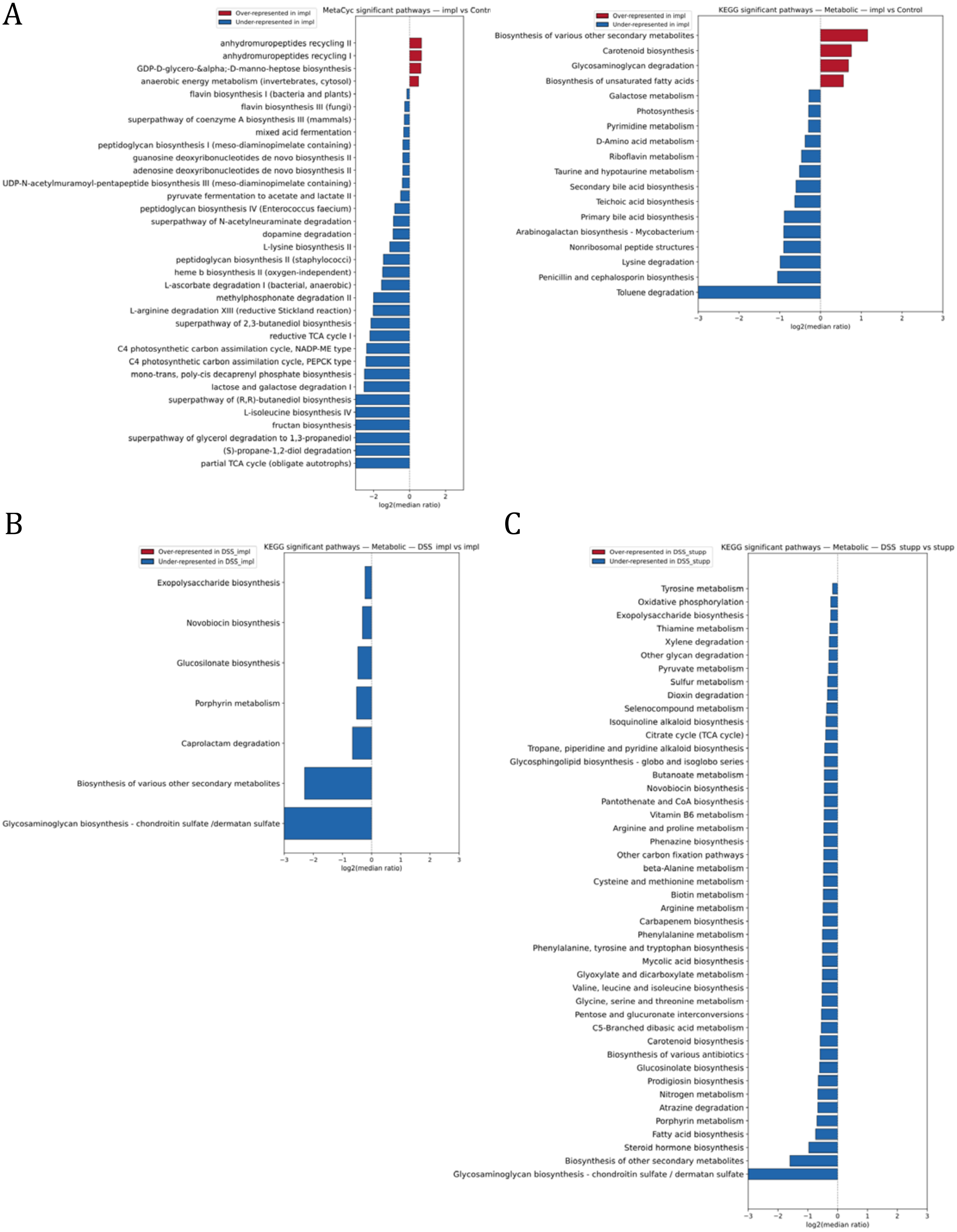


**Supplementary Figure 6**
